# Gene regulatory network determinants of rapid recall in human memory CD4+ T cells

**DOI:** 10.1101/2025.04.06.646912

**Authors:** Alexander Katko, Svetlana Korinfskaya, Anthony T. Bejjani, Seyifunmi M. Owoeye, Zi F. Yang, Akshata N. Rudrapatna, Sarah Potter, Joseph A. Wayman, Michael Kotliar, Leah C. Kottyan, Artem Barski, Emily R. Miraldi

## Abstract

Rapid recall is the hallmark of memory T cells. While naïve cells require days to mount effector responses to new threats, antigen-experienced memory cells produce cytokines within hours of repeat encounter. Memory establishment and the control of rapid recall across lifespan is poorly understood. Epigenetic poising is a likely mechanism. Compared to naïve, memory cells exhibit enhanced chromatin accessibility proximal to rapid recall genes, but the transcription factors (**TFs**) that establish, maintain and utilize these putative regulatory elements are unknown. We leverage single-nuclei (**sn**)multiome-seq (snRNA-seq and snATAC-seq) to characterize the dynamic activation responses of CD4+ T cell subsets and (2) reconstruct the underlying gene regulatory networks. Memory-associated TFs (MAF, PRDM1, RUNX2, SMAD3, KLF6) were predicted to orchestrate rapid recall. KLF6 binding to its predicted target genes was confirmed by ChIP-seq, while the memory-associated activities of all five factors replicated in independent scRNA-seq studies. Integrating GWAS data, we nominate CD4+ T cell populations and gene regulatory mechanisms that might underly genetic risk to immune-mediated diseases.

## Introduction

Memory CD4+ T cells are critical for protective immunity and tolerance. Upon initial antigen encounter, naïve CD4+ T cells undergo clonal expansion and, over several days, differentiate into cytokine-producing T helper (**Th**) cells. Depending on the cytokine and microenvironmental cues coincident with initial antigen encounter, the differentiating naïve cells acquire diverse immune capabilities: defense against intracellular pathogens (so-called Type 1 Th cells (**Th1**))(Mosmann et al., 1986), extracellular microbes (**Th17**)(Ivanov et al., 2009) and large extracellular parasites (**Th2**)(Sallusto, 2016; Van De Veerdonk & Netea, 2010), antibody class switching and affinity maturation (T follicular helper cells (**Tfh**))(Breitfeld et al., 2000; Johnston et al., 2009; Schaerli et al., 2000) and tolerance (induced regulatory T cells (**iTreg**))(Kanamori et al., 2016). The powerful defense capabilities of effector cells can cause collateral damage to the host. Most effector cells die following pathogen clearance, but some survive as memory cells(Hildeman et al., 2002; Reckling et al., 2008; Reinhardt et al., 2001). Memory cells possess a “rapid recall” ability to secrete cytokine and initiate clonal expansion within hours of re-exposure to antigen(Sprent & Surh, 2002). Rapid recall enables potent pathogen defense and is important for successful vaccination (Kaech et al., 2002). The phenomenon is remarkable because memory cells otherwise persist in a polar-opposite state (near quiescence) across decades and even lifespan (Hammarlund et al., 2003), their powerful effector and proliferation capabilities latent in the absence of antigen. Dysregulated rapid recall responses pose serious health threats: failure to mount contributes to chronic or life-threatening infections, while inappropriate, excessive responses drive allergic and autoimmune conditions(Aronica et al., 2004; Ishigame et al., 2013).

Despite its importance, the mechanisms underlying rapid recall and its long-term maintenance are poorly understood. Both naïve and memory CD4+ T cells can be activated by antigen-presenting cells (**APCs**). T cell receptor (**TCR**) stimulation by class II major histocompatibility (**MHCII**)-antigen complex and co-stimulation (e.g., via B7-CD28 axis) are sufficient for activation of both cell types. Memory cells have a higher barrier to activation than naïve cells(Mehlhop-Williams & Bevan, 2014) and are less dependent on co-stimulation. While differences in ZAP70, LAT and SLP76 have been described(Farber et al., 1997; Hussain et al., 2002; Kersh et al., 2003), the expression and activation kinetics of TCR signaling proteins are generally similar between naïve and memory T cells(W. Lai et al., 2011), and these signals converge to activate common downstream transcription factors (TFs), including AP-1(Macián et al., 2001), NFAT(Rao, 1994) and NFkB(Oeckinghaus & Ghosh, 2009; Teijaro et al., 2011). We and others thus hypothesized that the rapid recall ability of memory cells is regulated epigenetically, via local chromatin state that influences TF binding at key loci(Avni et al., 2002; Barski et al., 2017; Cuddapah et al., 2010; Fields et al., 2002). Initial evidence was provided by a study, which identified differential DNA methylation at cytokine gene promoters between naïve and memory cells, and epigenetic inheritance by daughter cells in the absence of TCR activation(Fitzpatrick et al., 1999).

Naïve T cell activation and effector cell differentiation dramatically alters the epigenome. Early single-gene and more recent ‘omics studies report alterations in DNA methylation(Durek et al., 2016; Hashimoto et al., 2013; Komori et al., 2015), deposition of lineage-specific histone marks at regulatory elements (Avni et al., 2002; Fields et al., 2002; Glasmacher et al., 2012; G. Wei et al., 2009), chromatin opening at key cytokine genes and changes in 3D chromatin structure(Ansel et al., 2006; Ciofani et al., 2012; Hatton et al., 2006; B. Lai et al., 2018; Lee & Rao, 2004; Messi et al., 2003). In resting memory T cells, histone modifications at regulatory elements of cytokine genes correlated with their rapid recall upon activation(Barski et al., 2017; Youngblood et al., 2013). These epigenetic modifications are thought to speed binding of activation-induced TFs to their regulatory elements, enabling faster gene expression kinetics in memory versus naïve T cells. Multiple TFs regulate expression of the so-called “rapid-recall genes”. These TFs include (1) TCR-inducible factors such as AP-1(Macián et al., 2001; Yukawa et al., 2020), NFATs(Rao, 1994) and NFkB(Sen & Baltimore, 1986), (2) cytokine-induced STATs(Darnell et al., 1994; Vahedi et al., 2012) and Th subset-specific regulators such as TBET(S. J. Szabo et al., 2000), GATA3, RORC(Hirose et al., 1994) or FOXP3(Fontenot et al., 2003; Zhou et al., 2008), and (3) non-inducible factors from ETS(WASYLYK et al., 1993) and RUNX(Bevington et al., 2016, 2017; Egawa et al., 2007; Gergen & Butler, 1988) families, posited to maintain the epigenetic state in resting memory T cells. Given the large number of potential regulators, a comprehensive approach is needed to elucidate the gene regulatory networks (**GRNs**) orchestrating rapid recall.

GRNs describe the control of gene expression by TFs. Genome-scale inference of GRNs from perturbation time-course gene expression and chromatin state data (e.g., using RNA-seq, ChIP-seq, ATAC-seq) provides a high-throughput means to infer thousands of regulatory interactions. CD4+ T cell biology has been an inspiring frontier for development of novel genome-scale GRN inference methods(Ciofani et al., 2012; N. Daga et al., 2024; Gustafsson et al., 2015; Henriksson et al., 2019; Miraldi et al., 2019; Wayman et al., 2023; Yosef et al., 2013). However, these prior works mainly focused on naïve cell activation/differentiation and not rapid recall specifically. Similarly, genomic characterizations (RNA-seq, ChIP-seq) of the rapid recall process, involving side-by-side comparison of TCR activation dynamics between memory and naïve cells, exist, but have not yet been integrated into a comprehensive GRN model. Their utilization for this purpose, however, is suboptimal, as these existing studies characterize epigenome at the population level, obscuring heterogeneity, an important GRN signal, in the human CD4+ T cell compartment.

Multi-modal, single-cell (**sc**)-genomics technologies provide a novel and necessary strategy for GRN modeling of rapid recall. Memory T cells are composed of populations of central (**TCM**) and effector (**TEM**) memory cells, distinguished by their homing, proliferation capacity and ability to produce cytokines(W. Lai et al., 2011; Sallusto et al., 1999). It has been notoriously difficult to resolve resting human TEM populations based on sc-transcriptomics alone(Cano-Gamez et al., 2020; P. A. Szabo et al., 2019). Thus, multi-modal, sc-technologies are needed.

Here, we apply single-nuclei (**sn**)-multiome-seq technology (parallel transcriptome and chromatin accessibility) to a T-cell activation time course of naïve and memory human CD4+ T cells, to elucidate the molecular drivers of rapid recall at genome scale. Multi-modal analyses (1) enabled resolution of the CD4+ T cell subsets across the time course, including resting TEM populations (Th1, Th2, Th17, Treg, cytotoxic and MHC class II-expressing populations) and (2) identified thousands of “rapid recall” genes and accessible chromatin regions exhibiting faster activation dynamics in memory versus naïve cells. Furthermore, hundreds of genes and thousands of accessibility regions distinguished resting memory and naïve populations; these molecular correlates represent potential drivers of rapid recall programming and maintenance at rest. We built genome-scale GRNs to identify TF regulators of gene expression and chromatin state across the CD4+ T cell populations, at rest and in response to T-cell activation. GRN analysis nominated a set of core TFs, RUNX2, PRDM1 (BLIMP1), MAF, SMAD3 and KLF6, responsible for both resting maintenance and early activation dynamics of rapid recall genes. Additionally, we shed light on TCM functional roles following activation, identifying potential for accelerated proliferation and acquisition of effector phenotypes. Integration of GWAS risk variants with our sc-resolved maps of chromatin accessibility implicated memory Th subsets and their rapid recall ability in genetic risk mediation of inflammatory and autoimmune diseases. Furthermore, we integrate genetic risk variants into the GRN to nominate disease-relevant regulatory mechanisms in CD4+ T cells.

## Results

### Resolution of naïve, TCM and TEM populations across a TCR activation time course for GRN inference

We designed T-cell activation time course experiments to unravel, at genome scale, the gene regulatory networks (**GRNs**) governing the rapid recall phenomenon in human memory CD4+ T cells (**Fig. 1A-C**). Given CD4+ T cell heterogeneity, we selected the single-nuclei (**sn**) multiome-seq technology to simultaneously characterize the chromatin landscape (accessibility by snATAC-seq) and gene expression (snRNA-seq) at single-cell resolution. CD4+ T cells from the blood of four healthy donors were prepared by negative magnetic selection. Within the CD4+ pool, the T effector memory (CD27^-^, **TEM**) population is low abundance but includes cells capable of high production of lineage-specific cytokines. Thus, we increased the fraction of TEM cells by partially depleting naive and T central memory (**TCM**) using anti-CD27 magnetic beads (**Methods**). Cells were then activated with anti-CD3/28 beads for early (2.5h), peak memory (5h) and late (15h) activation timepoints or remained resting and subjected to sn-multiome-seq analysis (16 experiments: 4 donors by 4 timepoints). To control for batch effects, resting cells from a fifth control donor were “spiked” into each experiment immediately before cell lysis; this strategy improved the quality of inferences in downstream analyses (**Fig. 1A, S1A-H, Methods**).

**Figure 1.**
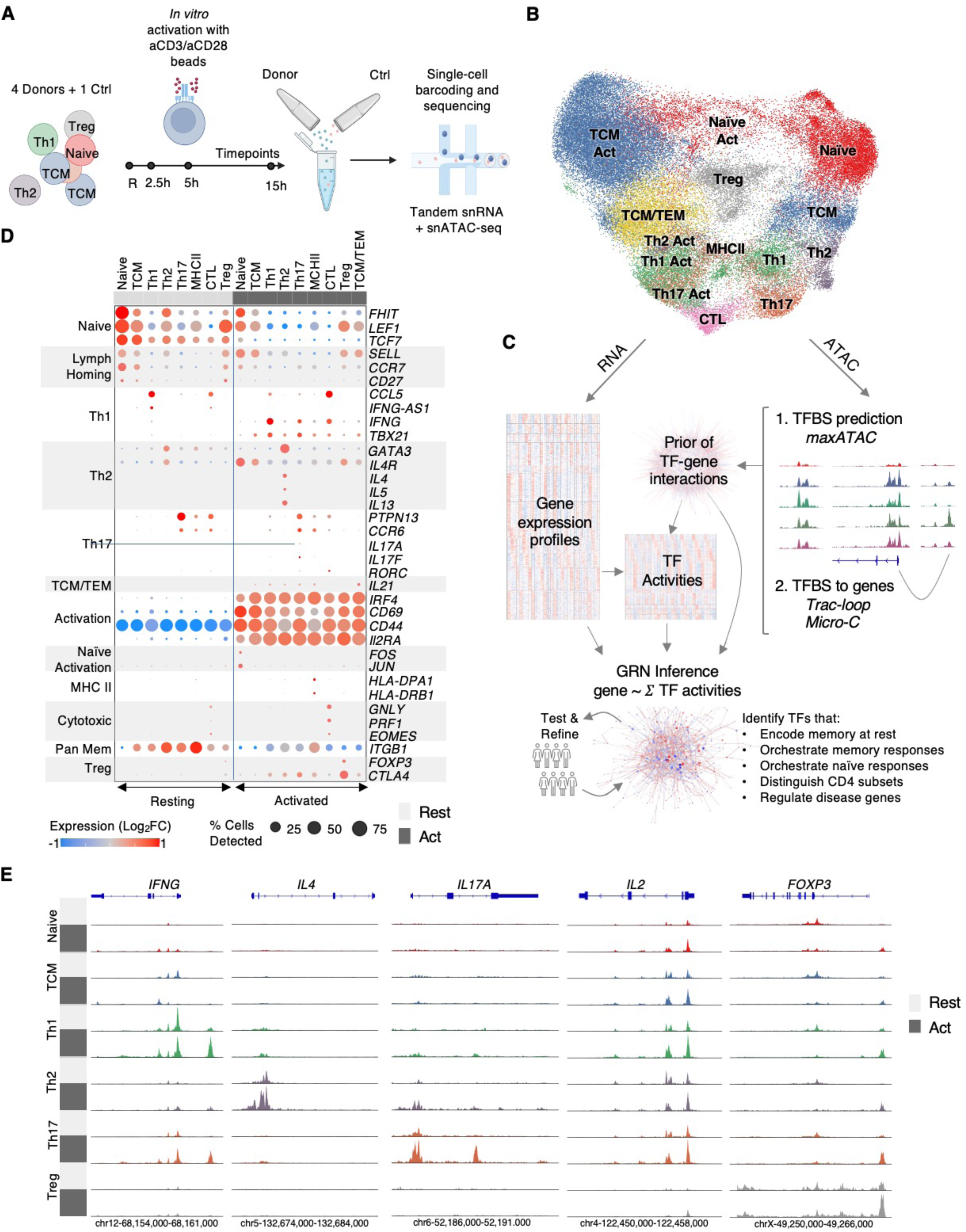
Resolution of naïve, TCM and TEM populations across resting and TCR-activation timepoints for GRN inference. **(A)** Experimental design: CD4+ T cells were activated using anti-CD3/CD28 beads and collected at four time points: resting, 2.5h, 5h, and 15h, for a total of 16 multiome-seq experiments (4 timepoints x 4 donors). To control for batch effects, cells from a fifth, “Ctrl”, donor (t=0) were spiked into each experiment immediately before lysis. (**B)** UMAP visualization, integrating transcriptome and chromatin accessibility from the resulting 65,862 high-quality nuclei. “**Act**” refers to activated cells from all activated timepoints (2.5-15h), combined. **(C)** Overview of the *Inferelator* for GRN inference from RNA-seq and ATAC-seq data. TF binding site (**TFBS**) prediction from ATAC-seq was accomplished using deep neural network models (**maxATAC**), when available. Putative TFBS were linked to target genes using experimentally derived promoter-enhancer interactions (Trac-loop, Micro-C) and linear proximity. The resulting “prior” of ATAC-derived TF-gene interactions enables estimation of protein TF activities and guides gene expression modeling (GRN inference). GRN predictions were tested and refined through analysis of external datasets. **(D)** Expression of select marker genes (transcripts per million (**TPM**), log_2_-normalized relative to the mean expression of each gene). **(E)** Normalized chromatin accessibility (see **Methods**) at select marker genes. (Panel A was created with biorender.com.)

The multiome-seq analyses yielded 65,862 high-quality CD4+ T cell nuclei, which composed 15 clusters (**Fig. 1B, Methods**), robustly recovered from each donor replicate (**Fig. S1I-L**). A cluster of “stressed” cells, mainly from 15h, lacked evidence of TCR activation, expressed stress genes and were removed from subsequent analyses (**Methods**). Next, we aggregated gene expression or accessibility signal from individual cells to generate pseudobulk profiles per CD4+ T cell subset (e.g., Th2), time point, and donor (**Methods**). Importantly, pseudobulk analyses mitigate data sparsity issues and leverage biological variation across replicates to more accurately estimate false discovery rate (**FDR**) and assess reproducibility(Heumos et al., 2023; Squair et al., 2021).

We used principal component analysis (**PCA**) to identify the major sources of variation in the gene expression and chromatin accessibility data. Activation was the dominant source of variation, principal component (**PC**) 1, in both data types (**Fig. S2A-F, Methods**). However, PCA also revealed data-modality differences. While PC1 was composed of genes peaking at 2.5-5h for transcriptome, the major trend (PC1) in the accessibility data corresponded to peaks monotonically gaining or losing accessibility over the time course. For gene expression, distinct temporal patterns dominated the gene expression data; they corresponded to PCs 1-3, which explained 44% of the data variance. As expected, PC1 had strong contributions from activation-dependent genes such as *IRF4* and *CD69.* For transcriptome, PC4 was the first PC to distinguish cell types: effectors (Th1, Th2, Th17, MHCII, CTL) from non-effectors (naïve, TCM, Treg). In contrast, for chromatin accessibility, the second most important trend, PC2, separated effectors from the non-effectors, while PC3 and PC4 captured a mix of cell-type and temporal variation. Hierarchical clustering complemented PCA. While transcriptomes formed four main clusters corresponding to timepoints (resting, 2.5h, 5h and 15h) (**Fig. S2G**), clustering of accessible chromatin profiles was driven by both cell type and time: all resting populations clustered together, while, for activation timepoints, effectors and non-effectors clustered separately (**Fig. S2H**). These results underscore the complementary information provided by snRNA- and snATAC-seq.

Although transcriptome alone distinguished activated naïve, TCM and TEM, chromatin accessibility was key to resolving (1) resting naïve versus TCM and (2) resting Th1, Th2 and Th17 populations (**Fig. 1D-E**). In the resting state, TCM and naïve cells share common signature genes (**Fig. 1D**), but accessible chromatin proximal to e.g., *IL2* and *CXCR5* (**Fig. 1E, Fig. S2I**), distinguished these populations at rest. Similarly, transcripts of signature cytokines and TFs were difficult to detect in resting Th1, Th2 and Th17 (**Fig. 1D**), but accessibility at these loci supported our annotations (**Fig. 1E**).

In addition to Th1, Th2 and Th17, we identified two small TEM populations, “MHCII”, “CTL”. Both populations expressed activation markers *IRF4* and *CD44* by 2.5h and lacked expression of homing TCM co-stimulatory molecule *CD27* (**Fig. 1C, S2J-K**). The “MHCII” population was distinguished by elevated expression of MHC Class II and antigen presentation genes (*CD74*, *CIITA*, *PSD4*), at baseline and upon activation (**Fig. 1C**, **S2L**).

The CTL populations expressed markers of cytotoxicity (*EOMES*, *GNLY*, *PRF1*), and activation further induced *GNLY* and *PRF1* expression. CTL also expressed Th1 and Th17 hallmark genes **(Fig. 1D).** Similar to Th1 and Th17 cells, *TBX21*, *IFNG* and *RORC* were only detected in CTL upon activation. Activated CTL expressed *RORC* as robustly as Th17 cells, but without concomitant *IL17A/F*.

Finally, the “TCM/TEM” population was uniquely detected at activation timepoints, with cell numbers increasing from 2.5 to 15h (**Fig. S1K-L**). The TCM/TEM cells expressed activation markers (early induction of *IRF4* and *CD44*, **Fig. S2I-J**), but with mixed central and effector memory features: expression of TCM markers *SELL* and *LEF1* and the cytokine *IL21* (**Fig. 1C**). In addition, TCM/TEM shared accessible chromatin regions with TEM populations at several effector gene loci (e.g., CCR family members). Globally, by both transcriptome and accessible chromatin, TCM/TEM clustered more closely with TCM than TEM, suggesting that TCM/TEM are a TCM population that gains TEM features upon TCR activation (**Fig. S2A-H**).

These clusters – and their associated transcriptomes and accessible chromatin profiles, resolved by donor and timepoint – served as input to the *Inferelator* algorithm for GRN inference (**Fig. 1C**)(Miraldi et al., 2019; Wayman et al., 2023). From the chromatin accessibility, we derived putative regulatory elements, using, for the first time, state-of-the-art neural network models *maxATAC* for TF binding site (**TFBS**) prediction(Cazares et al., 2023) and experimentally measured promoter-enhancer interactions(B. Lai et al., 2018; Oguchi et al., 2024) to associate predicted TFBS with potential target genes. The resulting prior of TF-gene interactions was used to estimate protein TF activities and guide gene expression modeling (GRN inference) (**Methods**)(Arrieta-Ortiz et al., 2015; Miraldi et al., 2019). The resulting GRN describes 123,314 regulatory (TF-gene) interactions between 428 TFs and 12,314 genes (**Table S1**). Analyses of the GRN nominated TFs involved in memory maintenance, activation responses in memory and naïve cells and subset-specific gene expression programs. To validate and refine the GRN predictions, we tested whether they (1) extrapolated to a substantially larger CD4 T cell activation sc-transcriptome dataset (655,349 cells, 119 human donors) (Soskic et al., 2022) and antigen-specific CD4 T cell activation contexts (Monian et al., 2022) and (2) were substantiated by TF ChIP-seq and perturbation data in CD4+ T cells.

### Thousands of rapid-recall genes exhibit faster, more extreme up- and down-regulation in activated memory cells

Having resolved the major CD4+ T cell populations, we investigated their transcriptional responses to T-cell activation. In total, we identified 10,042 dynamic, activation-dependent genes: 2086 unique to memory populations, 21 unique to naive cells and 7935 common to both (|log_2_(FC)| > 0.58, FDR=10% for at least one timepoint relative to t=0 in at least one population, (**Table S2**). We defined rapid recall (**RR**) genes as those with faster, more extreme dynamics in memory versus naive cells (**Methods**). This analysis identified 5831 RR genes (**Table S2**). The RR genes clustered according to dynamics (timepoint of peak expression, increasing or decreasing) and whether the trends were observed in the TEM populations (Th1, Th2, Th17, MHCII and/or CTL), the TCM population or both groups (**Fig. 2A-B**, **Table S2**). We functionally annotated the RR gene clusters and queried the GRN for predicted TF regulators (**Fig. 2C, S3A-E**, **Methods**, **Table S3**).

**Figure 2.**
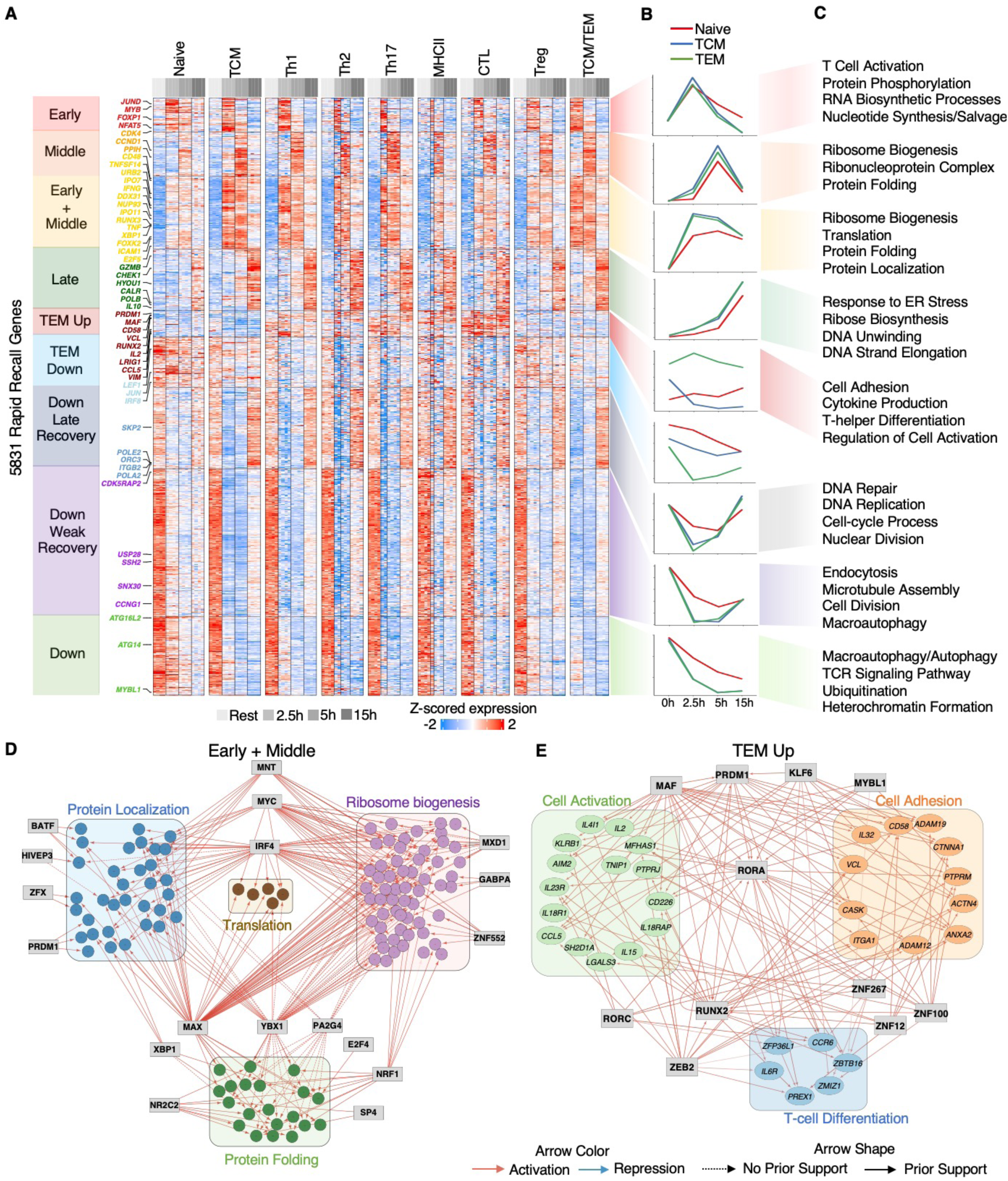
Rapid-recall transcriptional responses and their predicted TF regulators. **(A)** RR genes exhibited faster, higher-magnitude activation dynamics in memory compared to naive populations (n = 5831 genes, **Methods**). Donor, cell-type and timepoint-resolved pseudobulk gene expression values (VST counts) were z-scored and clustered. **(B)** Median gene expression profile per cluster, as a function of timepoint, for naïve, TCM and TEM populations; “TEM” represents the combined expression of Th1, Th2 and Th17 populations. **(C)** Select GO Biological Pathway enrichments per cluster (P_adj_ < 0.1). (No enrichments were identified for the TEM Down cluster.) GRN subnetworks for _select_ gene pathways enriched in the **(D)** “Early + Middle” and **(E)** “TEM Up” RR clusters. TFs are depicted as grey boxes and target genes as circles, colored according to their associated GO term. Target genes are limited to those (1) belonging to the given cluster and GO pathway and (2) regulated by at least one predicted TF regulator of the cluster (**Methods**). Solid arrows represent interactions supported by both gene expression modeling and the ATAC-derived “prior” network, while dashed arrows depict interactions supported by gene expression modeling alone.

TEM and TCM shared several upregulated gene clusters with maximal expression at early, middle, early and middle (“early+middle”) or late timepoints (332, 432, 715 and 696 genes, respectively). As swift cytokine production is a defining feature of rapid recall, early, “early+middle” and middle RR genes are enriched in pathways essential for cytokine production, including ribosome biogenesis, translation, protein folding and protein localization. For the early and middle RR genes, the GRN identified known (e.g., IRF4(Glasmacher et al., 2012), BATF(Sahoo et al., 2015), NFATC1(Clipstone & Crabtree, 1992; Jain et al., 1995) and NFAT5(Alberdi et al., 2017)) and novel (e.g., KLF6, POU2F2, HIVEP3) regulators in differentiation, cytokine response and immune effector function processes (Cluster C1,3,5,8, **Fig. S3D**, **Table S3**). The GRN newly identified TF regulators of protein synthesis and transport pathways with “early+middle” cluster dynamics (**Fig. 2D**). For example, YBX1 is a predicted regulator of ribosome biogenesis, protein folding and nuclear protein import, targeting ribosome assembly genes (*URB1*, *DDX46*), protein folding chaperones (*CCT5*, *CCT7*, *VAPA*), nucleoporins (*NUP93*, *NUP188*) and importins (*IPO11*, *IPO7*). The gene targets of IRF4, MYC and MAX are also enriched for ribosome biogenesis. BATF, HIVEP3, MAX, PRDM1 (BLIMP1) and MNT are predicted to regulate protein localization via induction of genes involved in intracellular transport (*COP1*, *XPOT*) and protein exocytosis (*CDC42*, *SCFD2*). Other TFs, including IRF4, MYC and NRF1, play roles in stress response, protein folding, biosynthetic processes (nucleotide, amino acid) and cell-cycle. Across the shared upregulated RR gene clusters with early and middle dynamics, many predicted regulators are themselves early (*NFAT5*, *NFATC1*, *BACH2*), early+middle (*CDC5L*, *GLIS3*) or middle (*XBP1*, *TBX21*, *IRF4*) rapid recall genes (Clusters T3, T4 and T5 in **Fig S3F-G**).

”Late” genes, maximal at 15h, include a second wave of genes to support massive protein synthesis (enriched in protein transport and folding, ER stress, cytokine production) and a new wave of genes involved in cell proliferation (enriched in DNA replication, cell cycle) (**Fig. 2A-C**). Many regulators (e.g., VDR, NFE2L1, PLAGL1, ATF5) of late-upregulated genes are themselves late-upregulated genes (Clusters T7, **Fig. S3F-G**).

Two gene clusters were specific to T effector memory: “TEM up” (260 genes) and “TEM down” (493 genes) (**Fig. 2A**). TEM-up genes are dramatically up-regulated (on average 3.4-fold) in TEM, relative to naive and TCM across all timepoints and display mixed dynamics, with most (87%) upregulated following activation. TEM-up genes are enriched for signaling pathways associated with canonical effector functions. The cluster includes cytokines (*IL2*, *IL22*), chemokine receptors (*CCR2*, *CCR5*, *CCR6*), TFs (*MAF*, *RORA*, *RUNX2*, *MYBL1*), and costimulatory molecules (*LIGHT*, *CD58*, *CD2*) (**Fig. 2C**). A core of mutually co-regulating TFs regulates these genes including KLF6, MAF, RUNX2, and others (**Fig 2E**). These TFs also exhibit high expression and GRN-predicted TF activity (**TFA**) in resting TEM (Cluster T5, **Fig. S3F-G**, **Methods**). Upon activation, transcription factor activities that are already high at rest are largely maintained, with a subset dynamically increasing (e.g., MAF, KLF6, PRDM1) or decreasing (e.g., SMAD3, RUNX2). The TEM-down genes are more lowly expressed in resting TEM and rapidly repressed further by 2h (**Fig. 2A-B**). In TCM and naïve, these genes are also downregulated, but to a lesser extent and more gradually. TEM-down genes include the naïve-associated TF *LEF1* and TCR signal transduction genes (*ZAP70*, *TEC*, *MALT1*).

Three rapid recall gene clusters exhibit more downregulation in memory versus naïve populations. In total, RR downregulated genes (2818) are nearly as frequent as upregulated genes (3003). The “down late recovery” cluster (793 genes) is comprised of genes repressed at the 2.5h and 5h timepoints, with return to or slightly exceeding baseline at 15h. Naïve cells exhibit similar dynamics but at lower amplitude. These genes are enriched for DNA repair, DNA replication, cell-cycle processes and nuclear division. They include DNA polymerases (*POLD3*, *POLE2*, *POLA2*), cyclin dependent kinases (*CDK2*), double-stranded break repair enzymes (*BRCA1*, *BRCA2*) and three of the six origin recognition complex genes (*ORC1*, *ORC3*, *ORC5*).

The “down weak recovery” cluster shows similar dynamics, but these repressed genes only recover to ∼50% of their baseline expression by 15h. They are involved in processes important for cell survival and homeostasis in the absence of antigen (e.g., endocytosis, macroautophagy). At 15h, these genes may regain expression to support a prolonged immune response by, for example, recycling organelles damaged during the activation. The gene cluster also contains *TOX*, which is involved in exhaustion and terminal differentiation(Alfei et al., 2019; Khan et al., 2019; Scott et al., 2019).

Finally, “down” genes monotonically decrease, with faster downregulation in memory cells. Some genes (*PTPRC*, *FYB1*, *PIK3CA*, *TRAT1*, *TNFRSF21*) are directly involved in TCR signaling. Others are connected to pathways that may interfere with activation: protein ubiquitination (*TRIM5*, *BIRC2*, *PRKN*), autophagy *(TOM1, SESN3*, *SIRT1*) and heterochromatin formation (*MECP2*, *PPM1D*).

While many genes exhibited RR dynamics across multiple memory populations, nearly half (45%, 2653 genes) were unique to a single memory subset (**Fig. S3A-C**). We thus sought to characterize the unique activation responses across the memory populations. Although individual genes were induced in the expected populations (*IL17A/F* in Th17 cells, *GNL* in CTL and antigen presentation genes in MHCII TEM), we detected very few pathway enrichments in the subset-specific rapid recall gene sets, indicating that the transcriptional responses, at a pathway level, are largely shared across TEM subsets.

In addition to the rapid recall genes, there were 4694 genes with similar activation dynamics in naive and memory cells (**Fig. S4A, Table S2**). These genes formed five clusters: early (816), middle (767), late (1067), down (759) and down late recovery (1285). Genes upregulated at early and middle timepoints were enriched for mRNA processing, splicing, translation, protein folding and other pathways supporting large-scale protein production. Their predicted regulators are IRFs, STATs, XBP1, KLFs and E2F4. Late upregulated genes were enriched for DNA repair and oxidative phosphorylation; predicted regulators included NR1H2, EGR3 and ATFs. Downregulated genes were involved in protein catabolism (“down”) and DNA repair and autophagy (“down recovery”). Notably, many RR regulators (PRDM1, MAF, RUNX2, FOXK2, KLF6, etc.) were not predicted to regulate genes with memory-naïve shared activation dynamics.

We examined whether the dynamic GRN predictions extrapolated to an independent CD4+ T cell activation scRNA-seq study, involving anti-CD3/28 stimulation at 0, 16, 40 and 120h. The zero and 16h timepoints enabled comparison with our first and last timepoints (0, 15h), for four CD4+ T cell populations (naïve, TCM, TEM and Treg, **Fig. S3H**). The TF activation dynamics were replicated in the larger study, with median Pearson correlation of 0.63 across the 188 TFs whose activity significantly changed during activation (**Fig. S4B**). Importantly, the *Soskic et al.* cohort (119 donors, 67 male, 52 female, over a range of ages: 19 to 79) enabled examination of TF dependence on sex and age. For the 378 TFs in our GRN that were also nominally expressed in the *Soskic et al.* dataset, time and cell type were the major sources of variation in the estimated TF activities. Limiting PCA to donor samples within a cell type-timepoint (e.g., TEM at 16h), age was a major source of TF variability only in naïve cells, at both 0h and 16h, as PC2 correlated significantly with age (ρ = 0.39 and ρ = 0.46, respectively, P_adj_ < 0.001). To identify individual TFs with sex- or age-dependence, we performed ANOVA and correlation analyses, respectively, for each TF (**Fig. S4C-D**). Mirroring the global analyses, 62 TFs exhibited age-dependent activities across multiple timepoints in naïve cells, but only one TF was similarly age-dependent in a memory cell type (TSHZ2 in TCM). The top-ranked TFs with age-increased activities were TSHZ2, PRDM1, KLF6, STAT3 and NFIX, while the top-ranked age-decreased TFs were HMGA1, ZEB2, BACH2, NFKB1 and PA2G4 (**Fig. S4D**). Among the 23 TFs exhibiting sex dependence, all were significantly sex-associated within the naïve compartment, while a subset were also sex-associated in memory cells (**Fig. S4C**). Only two TFs (HMGA1, ZBTB10) were elevated in males across the four populations (naïve, TCM, TEM, Treg), and ZNF496 was the only factor elevated in females, across the populations. In summary, analysis of the larger study enabled replication of predicted TF activity dynamics in a larger cohort, identified age as a significant variation source for TF activities within the naïve compartment only (62 out of 378 TFs) and a limited number of TFs with sex-dependent activities across cell types.

### Activated TCM diverge into highly proliferative and TEM-like cell states

Compared to TEM, TCM cells are more restricted to lymph nodes and exhibit an increased proliferative capacity and reduced production of effector cytokines(Lam et al., 2024; Sallusto et al., 1999). It is hypothesized that TCM serve as a reservoir for effector cells. Upon activation, murine TCM can differentiate into effector cells and migrate to peripheral tissues to orchestrate immune responses locally(Raphael et al., 2020; Wherry et al., 2003). In human cells, *in vitro* activation of sorted CD4+ TCM caused the TCM to acquire Tfh cytokine signatures(Rivino et al., 2004).

Our study provided an opportunity to explore human TCM function and regulatory mechanisms in the context of activation, relative to TEM. The TCM were composed of three populations: TCM, TCM/TEM (**Fig. 1B, C**), and “proliferating TCM”. The proliferating TCM population is a transcriptional state, apparent only through clustering of cell transcriptomes, independently of the accessible chromatin data (**Fig. 3A**). Indeed, we previously reported that proliferation genes are poised (i.e. have a chromatin environment similar to expressed genes) in resting T cells (Barski et al., 2009). Within our time course, the proliferating TCM state is primarily detected at 15h (95%) and accessed by both TCM and TCM/TEM cells (**Fig. 3A**). The three TCM populations were also reproducibly detected in the activated memory compartment of *Soskic et al.*(Soskic et al., 2022).

**Figure 3.**
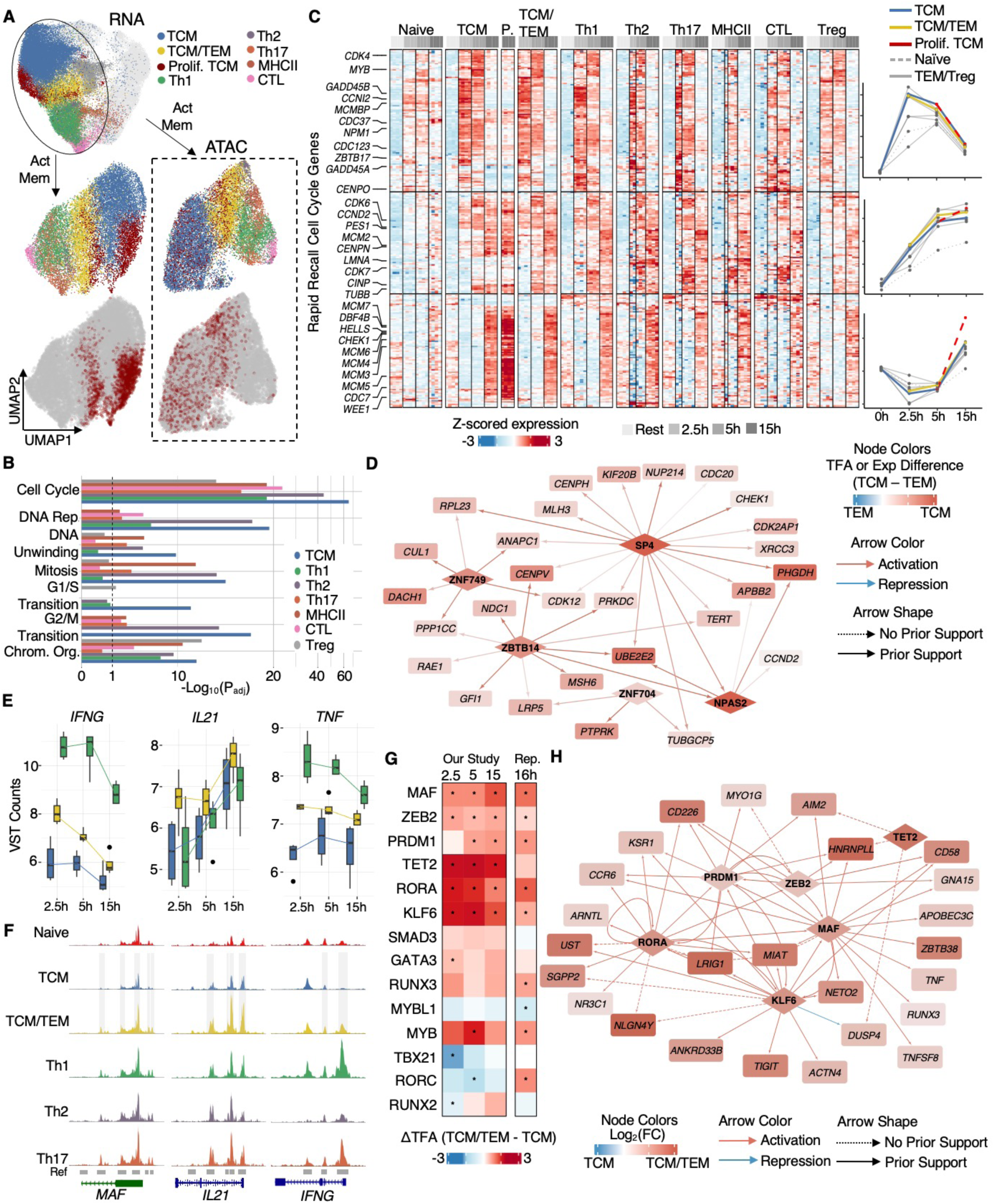
Characterization of the activated TCM compartment identifies proliferative and TEM potentials. **(A)** The proliferating TCM (“Prolif. TCM”) are a transcriptional state, identified via clustering of the transcriptome data alone, (RNA UMAP, upper left; light gray dots correspond to naïve or resting populations). UMAP visualization of activated memory cells by transcriptome alone (left lower panels) or accessibility alone (lower right panels) reveal that the proliferating TCM are a transcriptional state accessed by both TCM and TCM/TEM populations; upper panels show all activated memory populations, while lower panels highlight proliferating TCM, with other populations grayed out. **(B)** GSEA of each memory population’s 15h RR genes. The TCM population has the strongest enrichments in cell-cycle associated pathways. (Note TCM/TEM are excluded from this analysis, as they were not observed in resting conditions and rapid-recall genes are defined relative to the resting timepoint). **(C)** The gene expression dynamics of rapid-recall genes associated with cell-cycle. “P.” stands for “TCM-proliferating”. Line plots represent the average gene expression per population and cluster, as a function of time. **(D)** Gene regulatory network showing interactions between five TCM-associated TFs and their cell-cycle–associated target genes that are also elevated in activated TCM relative TEM. The five TFs were enriched for cell cycle gene targets. Node color indicates ΔTFA (TCM–TEM) for TFs and Δz-scored expression (TCM–TEM) for target genes. **(E)** TCM/TEM (yellow) have increased expression of several cytokines (*IFNG*, *IL21*, *TNF*) compared to TCM (blue); Th1 expression (green) is shown for reference. **(F)** Example loci where accessibility is elevated in activated TCM/TEM relative to TCM (2.5h, 5h, 15h combined); *P_adj_ < 0.1, FC > 1.5. **(G)** Core TCM/TEM TFs were prioritized based on differential TF activity between TCM/TEM and TCM (*P_adj_ < 0.1, t-test). **(H)** Core GRN for the TCM/TEM population, for select TFs and genes. Node color represents the expression fold change (genes) or ΔTFA (TFs) between TCM/TEM and TCM, averaged across the activation timepoints.

The proliferating TCM exhibit increased expression of DNA repair genes (*CHEK1*, *DNA2*) and DNA replication genes (*MCM10*, *POLQ* and *HELLS*) (**Fig. S4F**), suggesting enhanced proliferative potential. Relative to other memory populations, the induced RR genes of TCM at 15h are enriched for cell-proliferation genes: cell cycle, DNA replication, “G2/M transition” and others (**Fig. 3B, Table S4**). The GSEA results are supported by single-cell cell-cycle phase predictions (**Fig. S4G**). Proliferating TCM had the largest proportion of cells predicted to be in S, G2 or M phases, relative to all other cell-types at 15h (P_adj_ < 0.1, pairwise t-test).

We next visualized the dynamics of rapid recall genes involved in the cell cycle, identifying three major trends (**Fig. 3C**). The first gene cluster peaked at 2.5h in memory cells. The cluster includes *CDK4* and other regulators of early cell cycle progression: *MYB, MCMBP*, *CDC37* and *CDC123.* The second gene cluster increased over the time course; it included genes involved in G1 and S phase (*CDK6*, *CCND2*, *CINP*, *PES1*) and G2/M transition and mitosis (*CDK7, LMNA, TUBB,* CENP family). The third cluster consisted of late RR genes involved in DNA replication and S phase progression (*CDK2*, *CDC7*, *CHECK1*, *HELLS*, MCM and BRCA families). The proliferating TCM subset exhibited the highest expression of genes from this cluster, suggesting increased competency to start cell division.

Unbiased analysis of TF activity revealed a distinct regulatory program associated with the proliferation gene programs of TCM cells. Five TFs (SP4, ZNF749, ZBTB14, ZNF704 and NPAS2) were predicted to be more active in TCM than TEM cells, with target genes enriched in cell cycle genes (**Methods**, **Fig. 3D**). Some of these factors are associated with increased proliferation in cancer cells: SP4 in esophageal squamous carcinoma(L. Wei et al., 2024), ZNF704 in breast cancer(Yang et al., 2020) and uveal melanoma(Luo et al., 2023), and NPAS2 in acute myeloid leukemia cell lines(Song et al., 2019).

Cytokine-stimulated human TCM can be polarized into a “pre-TEM” state, suggesting that TCM might replenish TEM in inflammatory conditions(Geginat et al., 2001; Rivino et al., 2004). Here, our data suggests that T cell activation causes a subset of TCM cells to take on effector features, leading to the TCM/TEM population (**Fig. 1B, 3A**). The assertion that the TCM/TEM are derived from TCM is based on their overall similarity in both transcriptome and accessibility to TCM (**Fig. S2A-H)**. Relative to TCM, the TCM/TEM have increased expression of cytokine genes, including *TNF*, *IFNG*, *IL5*, *IL9,* and *IL21* (**Fig 3E**, **Fig. S4I**), and increased accessibility in these loci (**Fig. 3F**).

To nominate TFs contributing to the putative TCM transition to TEM, we created a core regulatory network for the TCM/TEM population (**Fig. 3G-H**, **Methods**). Several TFs core to TEM gene regulation (**Fig. 2G**), including MAF, KLF6, PRDM1 and RORA, were predicted to drive gene expression differences between TCM/TEM and TCM, suggesting that these TFs may also contribute to TCM acquisition of effector phenotypes. However, there were also important differences. The core TEM factor RUNX2 was notably absent, as *RUNX2* or its predicted activity was not increased in TCM/TEM-relative to TCM (**Fig. 3G, S4H**). Additionally, our TCM/TEM network highlighted a role for TET2, whose transcript was increased in TCM/TEM relative to both TEM and TCM populations (**Fig. S4F**). TET2’s DNA demethylation activity is crucial to reshaping the chromatin landscape during T helper differentiation(Baessler et al., 2023; Li et al., 2021). Thus, TET2 may likewise remodel chromatin to generate effector potential in TCM/TEM. In addition to the TCM/TEM-defining gene transcripts, most of the TF activities distinguishing TCM/TEM from TCM were reproduced by independent analysis of activated TCM from (Soskic et al., 2022)(**Fig. 3G, S4I**). Furthermore, the predicted transition of TCM to TCM/TEM is supported by a compositional increase in TCM/TEM cells from 16h to 40h (**Fig. S4E**).

Collectively, our findings suggest that activated TCM have greater proliferative potential than TEM and may serve as a reservoir for TEM cells. GRN analysis identified potential regulators of both processes.

### Most activation-responsive accessible chromatin regions exhibit rapid recall dynamics

Having characterized transcriptional responses to activation, we next investigated the underlying epigenetic landscape. Of the 324,074 accessible chromatin regions (“peaks”) identified in our study, 157,355 were dynamic (83,317 unique to memory, 5,452 unique to naive and 68,586 dynamic in both). Mirroring rapid recall genes, 46,134 “rapid recall peaks” were defined based on faster, higher-amplitude activation-induced opening or closing in memory versus naive cells (**Methods**). Unlike RR genes, which compose most dynamic genes (58%), RR peaks are a smaller fraction of all dynamic peaks (29%). In addition, while RR genes comprised nearly equal numbers of up- and downregulated genes, RR peaks primarily gained accessibility during activation (78%).

The RR peaks clustered into seven dynamic patterns (**Fig.4A-B**, **Table S5**). Accessibility in the “early open” cluster (4517 peaks) peaked at 2.5h and diminished slightly at subsequent timepoints. The “middle open” cluster (8363 peaks) monotonically gained accessibility throughout the time course. The “middle open TEM” cluster (6441 peaks) were elevated in resting TEM and gained accessibility across the time course (**Fig. 4B**). There were two “late-opening” peak clusters with maximal accessibility at 15h. They can be distinguished by the magnitude of accessibility increase between 0 and 15h, which was nearly 4-fold higher for the “late open strong” cluster (median log_2_(FC) = 3.2, 10,578 peaks) relative to the “late open weak” cluster (median log_2_(FC) = 1.3, 6097 peaks). Finally, activation rapidly diminished accessibility in the “rapid closed” (4651 peaks) and “rapid closed TEM” (5486 peaks) clusters. These peak clusters are exemplified by intronic peaks of several RR genes: *CD58* (early open), *VIM* (middle open), *MAF* (middle open TEM) and *LEF1* (rapid close) (**Fig. 4C**).

**Figure 4.**
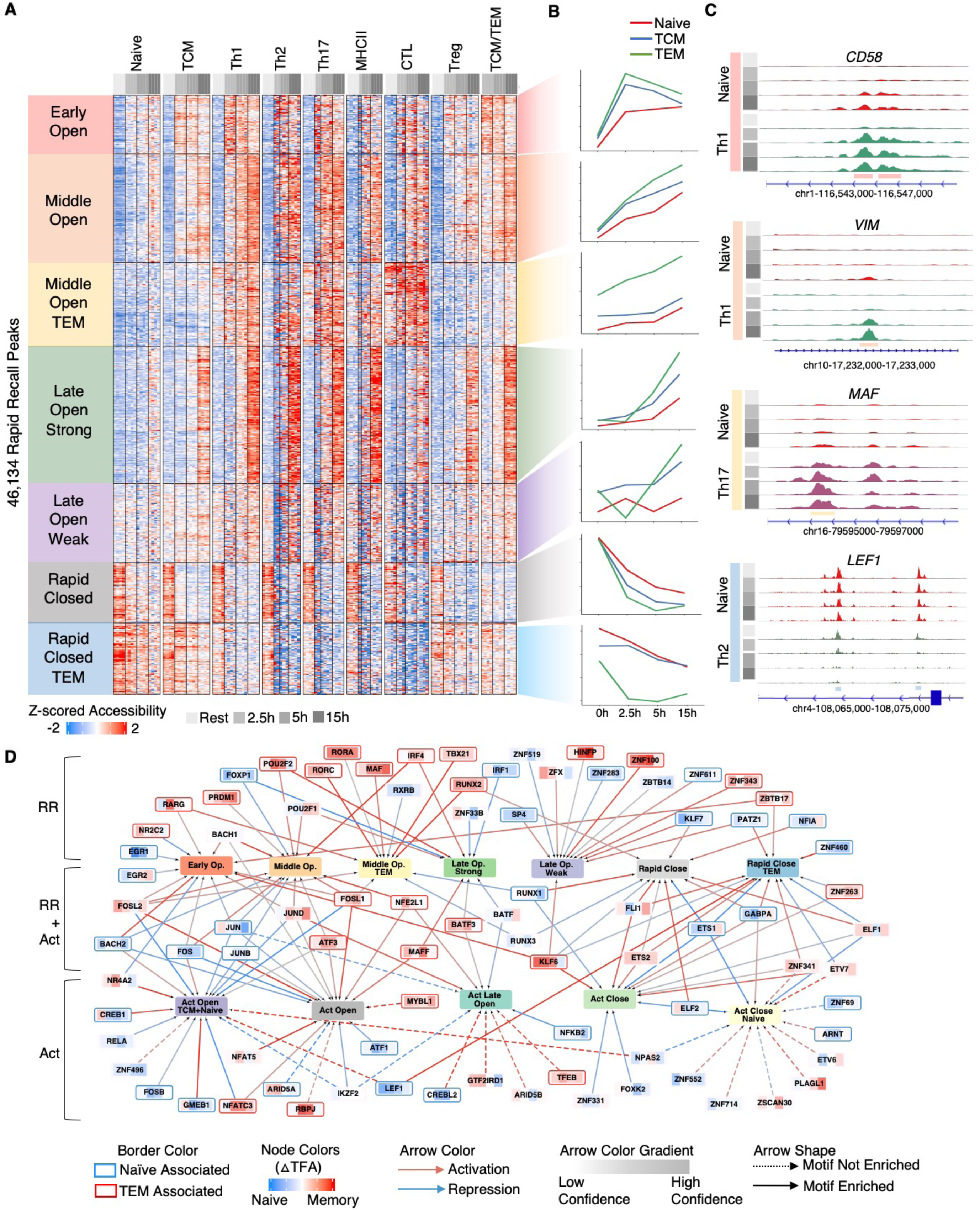
Chromatin accessibility rapid-recall dynamics and predicted TF drivers. **(A)** 46,134 accessible chromatin regions exhibit rapid recall dynamics (i.e., faster, higher-amplitude response to activation in memory than naïve cells). **(B)** The average accessibility signal per cluster, for naïve, TCM and TEM populations. (TEM combines Th1, Th2 and Th17 signal.) **(C)** Normalized ATAC signal at peaks with “early open” (*CD58*), “middle open” (*VIM*), “middle open TEM” (*MAF*) and “rapid closed TEM” (*LEF1*) dynamics. **(D)** A network model describing the TF regulation of activation-dependent chromatin accessibility. The network includes both RR peak clusters (Fig. 4A) and peaks with similar activation dynamics in memory and naïve populations (**Fig. S5A**). TF node colors indicate the difference in TFA between memory and naïve populations at each of the four timepoints; red outline indicates higher activity in memory cells, while blue indicates higher activity in naïve.

We also identified 39,622 activation-dependent peaks with similar dynamics between memory and naive cells, which we termed “shared activation-dependent peaks” or “act peaks” (**Fig. S5A**). These peaks clustered into five patterns. The “act open” cluster (8,136 peaks) exhibited early opening dynamics (maximal gain by 2.5h). The “act open TCM+naïve” cluster (7,927 peaks) similarly exhibited early opening dynamics, but the accessibility gain at 2.5h was at least two-fold higher in TCM and naïve populations relative to TEM. Across the CD4+ populations, the “act late open” cluster (8,733 peaks) steadily gained accessibility at each timepoint, achieving maximal accessibility at 15h. The “act close” cluster (8,691 peaks) closed in response to activation, with the greatest loss in accessibility between 0 and 2.5h. Finally, the “act close naïve” cluster (6,134 peaks) was composed of peaks with higher baseline accessibility that monotonically decreased upon activation in naive cells. Although few of the “act close naïve” peaks were dynamic in TCM, in TEM, accessibility was rapidly diminished by the 2.5h time point and then gradually increased to achieve accessibility on par with naïve cells at 15h.

We linked RR peaks (putative enhancers) with their potential gene targets. 53% of the RR peaks were promoter-proximal (+/-5kb of a gene TSS). Relative to all accessible chromatin regions in our study, RR peaks were more frequently gene-distal (OR=2.13, P<0.001) and more likely to form promoter-enhancer interactions, as detected from resting and activated CD4+ T cells (OR=1.44, P<0.001)(B. Lai et al., 2018; Oguchi et al., 2024). Similarly, activated peaks were also enriched for gene-distal regions (OR = 2.08, P<0.001) and promoter-enhancer interactions (OR = 1.39, p < 0.001).

We linked 55% of RR peaks and 54% of activation-shared peaks to potential target genes using Trac-loop and Micro-C interactions (28% and 27% peaks to distal genes, respectively) and a linear proximity rule (±5kb TSS). In general, RR peak sets were most enriched for the RR gene sets with corresponding dynamics (**Fig. S5B**). For example, peaks with early opening dynamics were most strongly enriched proximally to genes with “early”, “early+middle” and “TEM up” RR dynamics. The “late open strong” peaks were enriched proximally to the “late up”, “down late recovery”, “down weak recovery” and “down” RR genes, suggesting a role for these regions not only in promotion of rapid-up genes but also recovery of gene expression at 15h. Shared activation-dependent peaks were also strongly enriched proximal to both RR and activation-shared genes. For example, “act open” and “act open TCM+naïve” peaks sets were enriched proximally to early, middle and down RR genes as well activation-shared early up-regulated genes (**Fig. S5B**).

To identify TF drivers of activation-dependent chromatin remodeling (**Fig. 4A, S4A**), we built a novel multivariate modeling framework. While traditional TF motif enrichment analyses provided some clues about TF regulators of the accessibility patterns, motif enrichment analyses do not leverage the important relationship between TF activities and chromatin accessibility signal. Post-hoc assessment of this relationship is not trivial, when considering the multivariate, context-specific interactions of TFs across the 122 pseudobulk ATAC profiles (as shown in **Fig. 4A**). Thus, we explicitly modeled chromatin accessibility as a function of GRN-derived protein TF activities, using regularized regression to incorporate TFBS predictions (**Methods**). We built models for each of the twelve dynamic accessibility patterns (seven RR and five naïve-memory shared peak clusters), nominating a total of 84 TF regulators of chromatin dynamics (**Table S1**). Most of the TFs exhibited either TEM-associated activities (25 TFs, increased relative to naïve) or naïve-associated (30 TFs, **Methods**, **Fig. 4D**). Regulators of RR peaks were enriched for TEM-associated TFs (P_adj_ = 0.07, OR = 3.22), consistent with a role for memory-specific regulators in driving the accelerated chromatin dynamics characteristic of RR. Key TEM-associated regulators of RR peaks included PRDM1, RARG, RORA, RORC, MAF, IRF4, TBX21, RUNX2 and ZBTB17.

In contrast, TEM-associated TFs were underrepresented among regulators of the naïve-memory shared peak sets (P_adj_ = 0.03, OR = 0.34), suggesting distinct regulatory mechanisms for these genome regions. Indeed, among regulators of the shared dynamic peaks, there was an enrichment trend (P_adj_ = 0.15, OR = 2.14) for TFs lacking TEM- or naïve-associated activities, suggesting that shared activation peaks depend more on TFs broadly active across memory and naïve cells. Never-the-less, some TEM- or naïve-associated TFs were predicted to regulate shared regions of activation-dependent accessibility. For example, memory-associated MYBL1, NFATC3, RBPJ and ZNF263 were predicted to regulate shared activation peak sets, suggesting memory-specific regulatory mechanisms. Members of TF families (e.g., AP-1, ATF, ATF-like) were predicted regulators of shared activation-dependent peaks, with distinct family members associated with memory (BATF3, FOSL1, ATF3), naïve (FOSB, FOS, JUNB and JUN) or both cell types (BATF, FOSL2 and JUND). Naïve-associated TFs LEF1, KLF7, PATZ1, NFIA and ZNF460 were predicted to maintain accessibility of rapidly closing regions in naïve cells. Thus, the model highlights roles for memory- and naïve-associated as well as shared TF activities in regulation of activation-dependent chromatin accessibility dynamics.

In summary, hundreds of thousands of accessible chromatin regions exhibit activation-dependent dynamics and are linked to genes with corresponding dynamics. GRN modeling prioritized specific TFs, unique or common to naive and memory populations, responsible for orchestrating activation-induced chromatin dynamics in CD4+ T cells.

### Identification of TFs governing memory maintenance at rest

We previously hypothesized that the rapid recall ability of memory T cells is encoded in their resting-state epigenome(Barski et al., 2017). Other studies suggested that members of RUNX and ETS TF families are responsible for maintaining memory-defining open chromatin elements at rest. To uncover memory persistence mechanisms from our high-resolution multiome data, we identified accessibility chromatin regions differential between resting naive and memory populations and searched for candidate controllers: TFs (1) with binding site enrichment and (2) differentially expressed between resting memory and naiïve populations (**Fig. 5A-B**, **Tables S5, S6**). Our analysis yielded 32,661 candidate regulatory elements, potentially encoding T cell memory. The majority exhibited increased accessibility in memory populations (23,055 peaks, 71%), which were resolved into four clusters based on memory subset-specific patterns. The Th1/Th17/CTL cluster (4383 peaks) had the greatest accessibility in CTL followed by Th17 and Th1 populations, while the Th2/MHCII cluster (2904 peaks) was most accessible in Th2 and MHCII cells. These clusters were enriched for binding sites of their associated master regulators (TBX21, RORC, EOMES or GATA3) as well as regulators of the TEM activation response (PRDM1, MAF, RUNX2), nominated by GRN (**Fig. 2E**) and accessibility network (**Fig. 4D**) analyses. These TFs were also supported by elevated transcript in the corresponding memory populations. The other memory-associated clusters were distinguished by relatively low (13,860 peaks) or high (2036 peaks) accessibility in Treg. By motif enrichment and transcript expression, BATF, MAF and RUNX2 are implicated in the resting maintenance of both clusters. Finally, two clusters exhibited increased accessibility in naive cells, were moderately accessible in TCM and distinguished by moderate or low Treg accessibility (“naive+Treg” and “naive” clusters, respectively). Motif enrichment and gene expression criteria nominated several TFs, including four LEF1, ETS1, TCF7 and EGR1, as regulators of the naïve-associated clusters.

**Figure 5.**
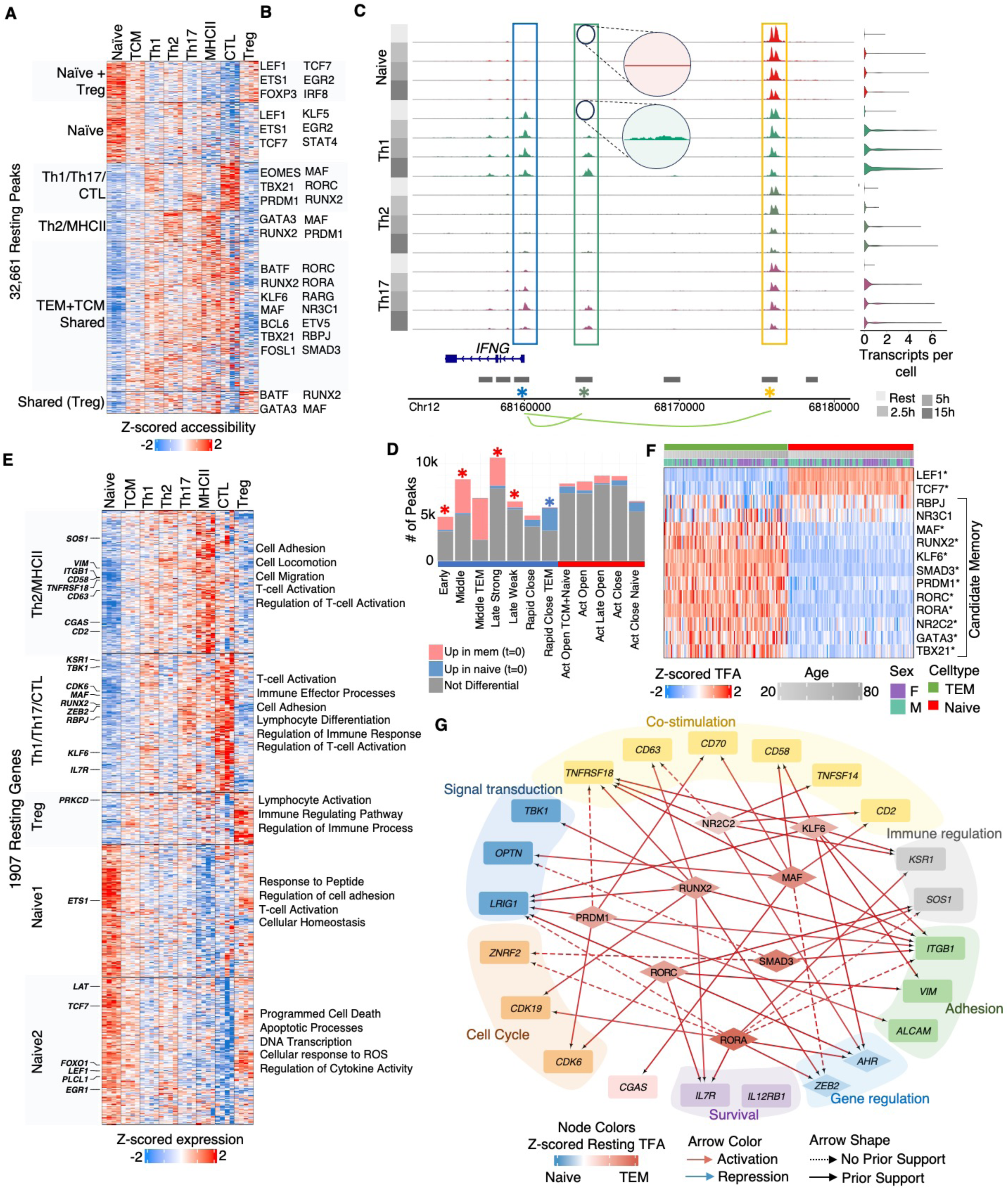
TF regulation of memory-associated accessible chromatin regions and genes at rest. **(A)** Clustering of chromatin accessibility regions differential between naive and at least one memory population at rest (n = 32,219 peaks). **(B)** Putative TF regulators per accessibility cluster in (A), where TFs (1) are differentially expressed between resting naive and memory populations, with a fold-change that positively correlates with the cluster’s accessibility signal, and (2) have motif enrichment in the corresponding peak cluster. **(C)** Normalized ATAC signal at the *IFNG* locus for naïve, Th1, Th2 and Th17 cells. The blue box highlights a RR peak at the *IFNG* promoter, with accessibility in Th1 and, to a lesser extent, Th17 cells. Green and gold boxes highlight accessibility in *IFNG* distal regulatory regions (supported by Trac-loop); these peaks show opposite trends: RR opening (green box) and RR closing (gold box). Zoom-in boxes show accessibility in resting Th1 but not naive populations. Violin plots display *IFNG* expression. **(D**) For each RR or memory-naïve shared activation peak set (Fig. 4A**, S5A**), the number of peaks that were also differential in resting memory and naïve cells (A) is indicated. Red (or blue) asterisk indicates significant enrichment of “memory-up” (or “naïve-up”) peak sets by Fisher exact test, P_adj_ < 0.1. **(E)** 1907 genes were differentially expressed between naïve and at least one memory population at rest. (F) For memory-encoding TFs nominated by our study, TF activities are verified in resting TEM and naïve cell populations from *Soskic et al*. (n = 119 donors); naïve-associated LEF1 and TCF7 are shown for reference. Asterisk indicates memory-dependent TFA (P_adj_ < 0.1, t-test) in *Soskic et al*. Donors are ordered according to age, with sex indicated. **(G)** GRN subnetwork of “resting memory maintenance”, highlighting TFs and target genes in resting memory cells predicted to contribute to rapid recall. Target genes are colored by function.

We next explored how regions of resting memory-associated chromatin might underlie rapid recall dynamics of peaks and genes. The phenomenon is exemplified by the *IFNG* locus, in which a rapid recall peak, linked to the *IFNG* by Trac-loop, exhibited low-level but significantly increased accessibility in Th1 relative to naïve cells prior to activation (green box, **Fig. 5C**). Indeed, many of the rapid recall peaks were differentially accessible in resting memory versus naive cells. Apart from the “late weak open” cluster, all other RR peaks with opening dynamics were significantly enriched for peaks open in resting memory versus naive cells (**Fig. 5D**), supporting that pre-existing chromatin accessibility in resting memory cells contributes to rapid recall. Likewise, the RR peak cluster with closing dynamics (Close TEM) was enriched for peaks with less accessibility in resting memory versus naïve cells, suggesting epigenetic mechanisms for faster, RR closing of accessible chromatin regions. In contrast, none of the activation-dependent peak sets with shared memory-naïve dynamics were enriched for resting memory-associated accessible chromatin (**Fig. 5D**).

Prior studies found little evidence that rapid recall was driven by enhanced expression or activity of canonical TCR signaling proteins(W. Lai et al., 2011). Here, we reexamine this possibility, taking advantage of sc-resolved memory subset transcriptomes to discover gene expression differences between memory and naïve populations at rest that may contribute to rapid recall. In total, there were 1907 genes differentially expressed between naïve and at least one memory population (1028 upregulated in memory, 869 upregulated in naïve) (**Fig. 5E, Table S2**). Using the Trac-loop and Micro-C, resting memory-up peaks were enriched for enhancer links to resting memory-up genes, while resting naive-up peaks were enriched for resting naive-up genes (one-sided Fisher’s exact tests, P < 0.01). GSEA revealed a common theme of cell adhesion, activation and migration among the upregulated memory genes. The “Treg” memory-associated gene cluster (highest resting expression in Treg, 161 genes) was enriched for immune regulation pathways and included Treg-associated genes like *FOXP3*, *CTLA4* and *IKZF2*. Naïve-upregulated genes included TCR signaling molecules (*LAT*, *LCK*, *PLCL1*) and naive-associated (*FOXO1*, *LEF1*, *TCF7*) and other TFs (*ETS1*, *EGR1*).

Using the GRN, we prioritized TF regulators of the rapid recall potential at rest (from **Fig. 5B**), based on regulation of the resting memory-associated genes (**Fig. 5E**, **Methods**). These candidates were further refined by interrogation of the *Soskic* et al. dataset, in which all but two of the twelve predicted resting-memory core TFs (RBPJ, NR3C1) were supported in the larger cohort (**Fig. 5F**). TFs KLF6, MAF, RUNX2, NR2C2, SMAD3, PRDM1, RORA and RORC were broadly active across multiple resting TEM populations (**Fig. 5G**). TF regulation of co-stimulatory molecules *CD58*, *CD2*, *TNFRSF18*, *CD63*, *TNFSF14* and *CD70* at rest may contribute to RR. For example, CD58 and its binding partner CD2 increase T-cell responsiveness to activation and downstream cytokine production through PLCγ1 dependent Ca^2+^ signaling(S. Q. J. Liu & Golan, 1999); thus, baseline increase in *CD58* and *CD2* expression may drive rapid-recall responses *in vivo*. Collectively, our analyses nominate accessible chromatin regions, transcripts, TFs and GRNs, uniquely utilized by resting memory cells to enable their rapid recall ability.

### Memory-associated TFs are implicated in early differentiation of naïve cells to effectors

Limiting to naïve cells, we used pseudotemporal ordering to characterize the naive compartment’s heterogeneity across resting and activated timepoints. The pseudotemporal ordering correlated with the temporal dynamics of our design, excepting some unstimulated cells (low *IRF4*, *CD44* expression) from all timepoints concentrated early in the trajectory (**Fig. 6A-B**). Next, we visualized module scores summarizing accessibility in the putative “memory-encoding” peak sets (**Fig. 5A**) on a per-cell basis (**Fig. 6C-E**). A subset of naive cells at late pseudotime, mostly from 15h, gain accessibility in regions open in resting memory and lose accessibility in regions defining resting naïve cells. These so-called “precocious” naive cells acquire memory-associated chromatin features faster than other activated naïve cells at 15h. This trend extends beyond the resting memory-specific regions to include accessibility changes in the RR and shared activation peak sets. Relative to other naïve activated cells at 15h, the precocious naive gained (or lost) more accessibility in peak sets opening (or closing) in activated cells (**Fig. 6C-D**).

**Figure 6.**
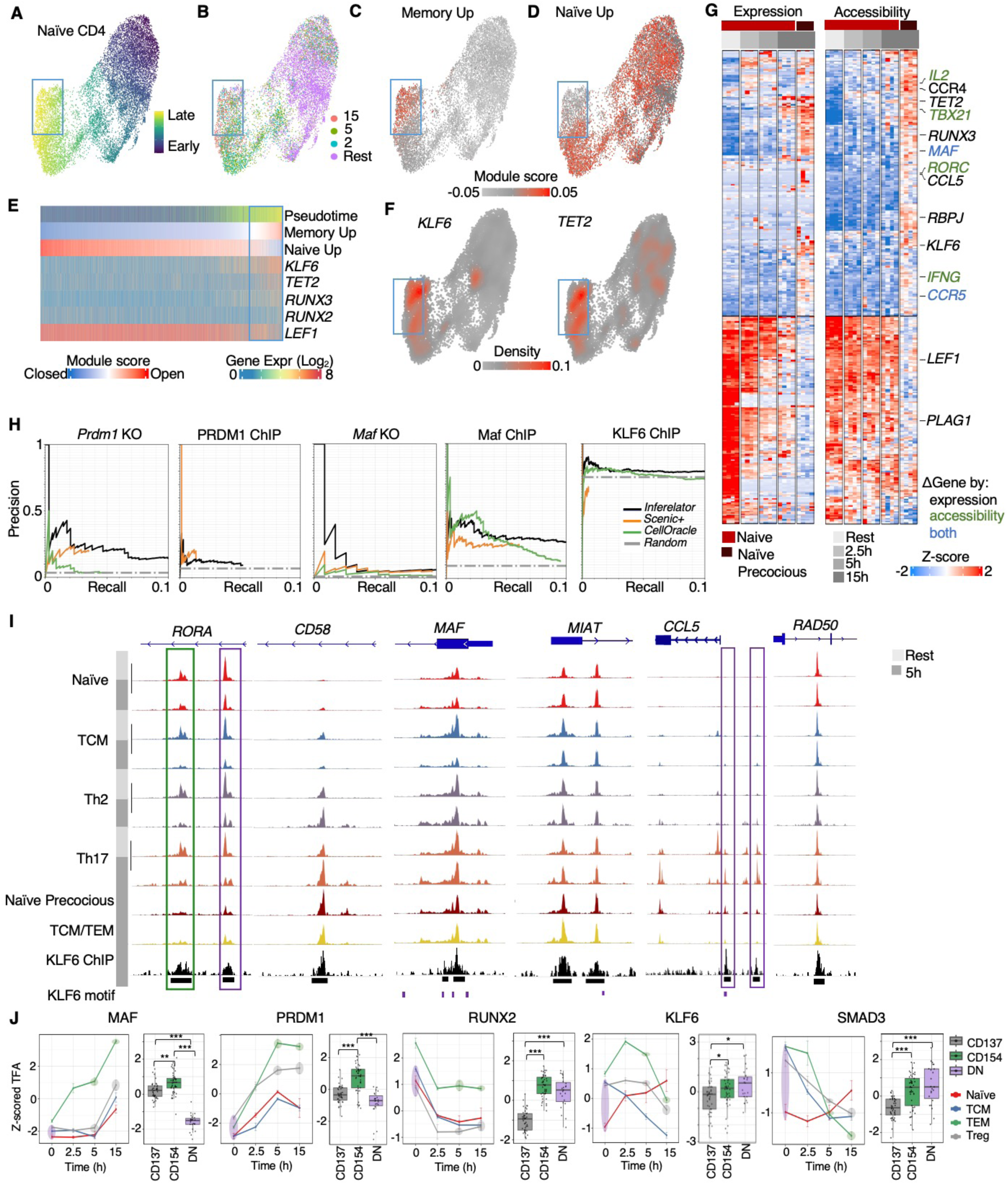
Enhanced activation kinetics in a naïve-cell subset and experimental testing of GRN predictions. WNN UMAP of all naïve CD4+ T cells colored by **(A)** predicted pseudotime, **(B)** experimental timepoint and **(C, D)** average accessibility of peaks from either the naïve-associated or memory-associated peaks at rest (from Fig. 5A, **Methods**). Scores > 0 indicate above-average peak-set accessibility, relative to all other CD4+ T cells. A subset of late pseudotime “precocious” naïve cells (blue box) shows positive “Memory Up” scores. **(E)** Per-cell “Memory Up” and “Naïve Up” accessibility scores and expression (log_2_(TPM)) of select TFs across naïve-cell pseudotime. **(F)** Density plots of TF expression in naïve cells. **(G)** Co-clustering of gene expression and summed proximal accessibility for 849 genes with differential expression or proximal accessibility between precocious naïve (defined as “Memory Up” module score > 0.025) and other activated naïve cells at 15h (P_adj_ < 0.1). **(H)** Precision–recall performance of our GRN (*Inferelator*) on prediction of experimentally supported regulatory interactions for MAF, PRDM1 and KLF6 in CD4+ T cell datasets. The total number of “gold-standard” regulatory interactions were: 1262 (*Prdm1* KO), 1362 (PRDM1 ChIP-seq), 636 (*Maf* KO), 1159 (*Maf* ChIP) and 11,439 (KLF6 ChIP-seq). GRNs derived from other methods are shown for reference. **(I)** KLF6 ChIP-seq linear fold-change enrichment signal and multiome-derived ATAC-seq signal profiles (counts per million) at KLF6 ChIP-seq peaks. KLF6 motifs are indicated. Genomic coordinates are: *RORA*: chr15:60,830,000-60,850,000, *CD58*: chr1:116,560,000-116,580,000, *MAF*: chr16:79,595,000-79,605,000, *MIAT*: chr22:26,640,000-26,648,000, *CCL5*: chr17:35,870,000-35,905,000, *RAD50*: chr5:132,555,000-132,558,000. **(J)** A comparison of TF activities for the five memory-associated core TFs were compared between our aCD3/CD28 activation time course (left panel) and antigen-specific activation (right panel), using scRNA-seq data from *Monian et al.*, in which T cells from peanut-allergic individuals were stimulated *ex vivo* with peanut antigen for 20h and sorted into CD137⁺ (enriched for peanut-specific Treg), CD154⁺ (peanut-specific TEM), and double-negative (**DN**) populations (n = 109 donor timepoints). Population-dependent activities were determined by t-test (P_adj_ < 0.1 = *, < 0.001 = **, < 0.00001 = ***). In our data (left panel), color-coded ovals denote approximate cell-type-timepoints aligning with the peanut-stimulation populations, accounting for antigen processing and presentation time (e.g., CD154⁺ at 20h corresponds to ∼5–15h TEM). See text for more details.

We explored potential TF regulators of the precocious naive cells (**Fig. 6E-F**). Like resting and activated TEM, precocious naïve cells exhibited increased *KLF6* expression (**Fig. 6E-F**). Additionally, naive precocious displayed increased expression of TEM regulators *RUNX3* and *TET2*(Cruz-Guilloty et al., 2009; Ichiyama et al., 2015). Genome-wide, hundreds of genes have significant differences in either expression (667) or proximal accessibility (397) between naïve precocious and other 15h naive activated cells (**Fig. 6G**, **Table S7**). Some effector (*MAF*, *CCR5*) and chromatin remodeling (*TET2, RORC, TBX21*) genes were already elevated in precocious naïve, while other genes (*IL2*, *IFNG*, *RORC* and *TBX21*) had a significant increase in proximal accessibility but not expression, suggesting transcriptional priming.

Thus, the putative memory-encoding peaks: (1) exhibit accessibility in resting memory but not naive cells and (2) gain accessibility in activated naive cells by 15h. As a result, the “memory-encoding” peaks appear functionally important to both rapid recall (memory cells) and the differentiation of activated naive cells into effectors. Both processes implicate shared TFs (*KLF6*, *MAF*) as well as unique TFs (*RUNX2* unique to memory and *TET2* and *RUNX3* unique to precocious naive cells).

### Testing of memory-associated TF regulatory interactions and their implication in antigen-specific activation

We generated ChIP-seq data to validate predicted TF-gene interaction for KLF6, a novel TF regulator of resting memory maintenance, rapid recall and the acquisition of effector memory transcriptional signatures in activated TCM and naïve populations. In activated CD4 T cells, we detected thousands of KLF6 binding sites, with many proximal to predicted KLF6 target genes (**Methods**). Integration of the highly-resolved chromatin accessibility profiles provided insights into the dynamics and cell-type specificity of TF binding at KLF6 binding sites in CD4+ T cells (**Fig. 6I**). In the *RORA* locus, KLF6 bound both a rapid-recall opening and a RR closing region (purple and green box, respectively). Accessibility at KLF6 binding sites in *CD58, MIAT and MAF* was already elevated in resting memory cells, with activation-induced accessibility gain in precocious naïve and TCM/TEM populations. KLF6 binding sites at the *CCL5* locus exhibited subset-specific increases in accessibility following activation in Th17 and precocious naïve cells (purple boxes). In contrast, the KLF6 binding site in *RAD50*, within the Th2 cytokine locus, was accessible across CD4+ T cell populations, in both resting and activation contexts.

To evaluate the quality of predicted TF-gene regulatory interactions at genome scale, we used our KLF6 ChIP-seq, along with publicly available ChIP-seq and TF knockout (**KO**) data for two other GRN-predicted, memory-core TFs MAF and PRDM1 (Guo et al., 2022; Wayman et al., 2023) (**Methods**). Following the standard in the field (Marbach et al., 2012), we evaluated precision (fraction of GRN TF-gene interactions supported by KO or ChIP-seq data) and recall (fraction of KO or ChIP-supported interactions recovered by the GRN) as a function of GRN interaction confidence scores. For reference, we compared the performance of our method (*Inferelator*) to other popular GRN inference methods(Bravo González-Blas et al., 2023; Kamimoto et al., 2023), demonstrating on-par or superior predictive performance for these three factors (**Fig. 6H**).

The predicted TF dynamics and their association with specific CD4+ T cell populations at rest or upon activation, derived from sn-multiome analysis of four donors, were first replicated using scRNA-seq of a similar anti-CD3/CD28 activation time course across a much larger cohort (119 donors) (**Fig. S3B**,**H**, **S4E-F**,**I**, **4G**, **5F**). Next, we examined whether the GRN predictions were supported in the more physiological context of antigen-specific T cell activation. For the comparative analysis, we estimated TF activities from scRNA-seq of peanut-activated CD4+ T cells, isolated from peanut-allergic individuals undergoing peanut oral immunotherapy (n = 12 donors)(Monian et al., 2022). In their design, peripheral blood mononuclear cells (PBMCs) were isolated, incubated with peanut antigen for 20h and activated CD4^+^ TEM and Treg were enriched based on CD154 and CD137 induction, respectively. CD4^+^ T cells, double-negative (**DN**) for these markers (CD154^-^, CD137^-^), served as a control. Accounting for antigen processing and presentation time in their design, we reasoned that the activated TEM and Treg in their dataset roughly corresponded to the 5 and 15h aCD3/CD28 activation timepoints in our dataset. As TCM and naïve cells compose the majority of peripheral CD4+ T cells, we compared TF activities from their DN timepoint to the average activities of resting naïve and TCM in our dataset.

For the memory core (KLF6, MAF, PRDM1, RUNX2, SMAD3), there was qualitatively good agreement between the predicted TF activities of activated TEM and Treg and resting DN cells between antigen-specific (peanut) and aCD3/CD28 activation (**Fig. 6J**). For example, our analysis predicted that the activities of MAF and PRDM1 would be highest in activated TEM, followed by activated Treg and lowest in resting DN cells. Similarly, we predicted that RUNX2 activity would be lowest in activated Treg and highest in resting DN cells. For KLF6, our dataset predicted that the three populations would have overlapping ranges of activity, and, for SMAD3, our analysis predicted slightly elevated activity in DN. Both trends were consistent with the antigen-specific TF activity estimates. We next undertook an unbiased comparison of TF activities between the datasets, using ANOVA to identify TFs whose activities were not static across the three groups (activated Treg, activated TEM and resting DN cells), in the peanut study or as simulated from our dataset (**Methods**). Side-by-side comparison of the resulting 90 TFs confirmed agreement between the predicted TF dynamics in antigen-specific and aCD3/CD28 contexts for most TFs (64 TFs, **Fig. S6**). As anticipated, there was also some disagreement, due to expected differences in the underlying molecular biology and/or our crude process of simulating expected TF activities for CD137^+^, CD154^+^ and DN CD4^+^ populations from our dataset. Disagreement in TF activity predictions were frequently driven by more subtle differences in the relative ranking of the activated memory populations (Treg and TEM), whereas larger differences distinguishing activated memory and resting DN populations were typically captured (**Fig. S6**). Although further benchmarking is needed, these results suggest that the GRN could be used to identify activated CD4 T cells (Treg or TEM) from physiological scRNA-seq studies, in the absence of cell sorting strategies for activated cells.

Based on reproducibility of GRN predictions in external cohorts and orthogonal evaluation of TF-gene interactions by TF perturbation and ChIP-seq data, we were emboldened to use the GRN to identify potential molecular mechanisms for human genetic risk variants in CD4+ T cells.

### Memory-associated accessible chromatin regions and subset-defining GRNs are implicated in mediating genetic risk for allergic, autoimmune and inflammatory diseases

CD4+ T cells and their activation states mediate genetic risk for autoimmune and inflammatory diseases(Calderon et al., 2019; K. Daga et al., 2024; Harley et al., 2018; Soskic et al., 2022; Wayman et al., 2024). Genetic risk variants for these and other complex diseases are predominantly noncoding, enriched in regulatory elements and, if causal, thought to alter TF binding and gene expression(Farh et al., 2015). To examine whether accessible chromatin regions defined by our study (e.g., rapid recall-associated) might mediate genetic disease risk, we utilized the Regulatory Element Locus Intersection (**RELI**) algorithm(Harley et al., 2018) to systematically assess the colocalization of GWAS-identified sets of genetic risk variants with accessible chromatin regions (**Table S8**, **Methods**). Of the 387 disease risk variant sets (phenotypes) tested, 44 significantly colocalized with at least one of our 18 accessible chromatin maps (FDR = 10%, **Fig. 7A**). Accessible chromatin regions distinguishing resting memory from naïve populations were most significantly enriched for genetic risk variants, followed by accessible chromatin regions that opened upon activation, both RR and memory-naïve shared.

**Figure 7.**
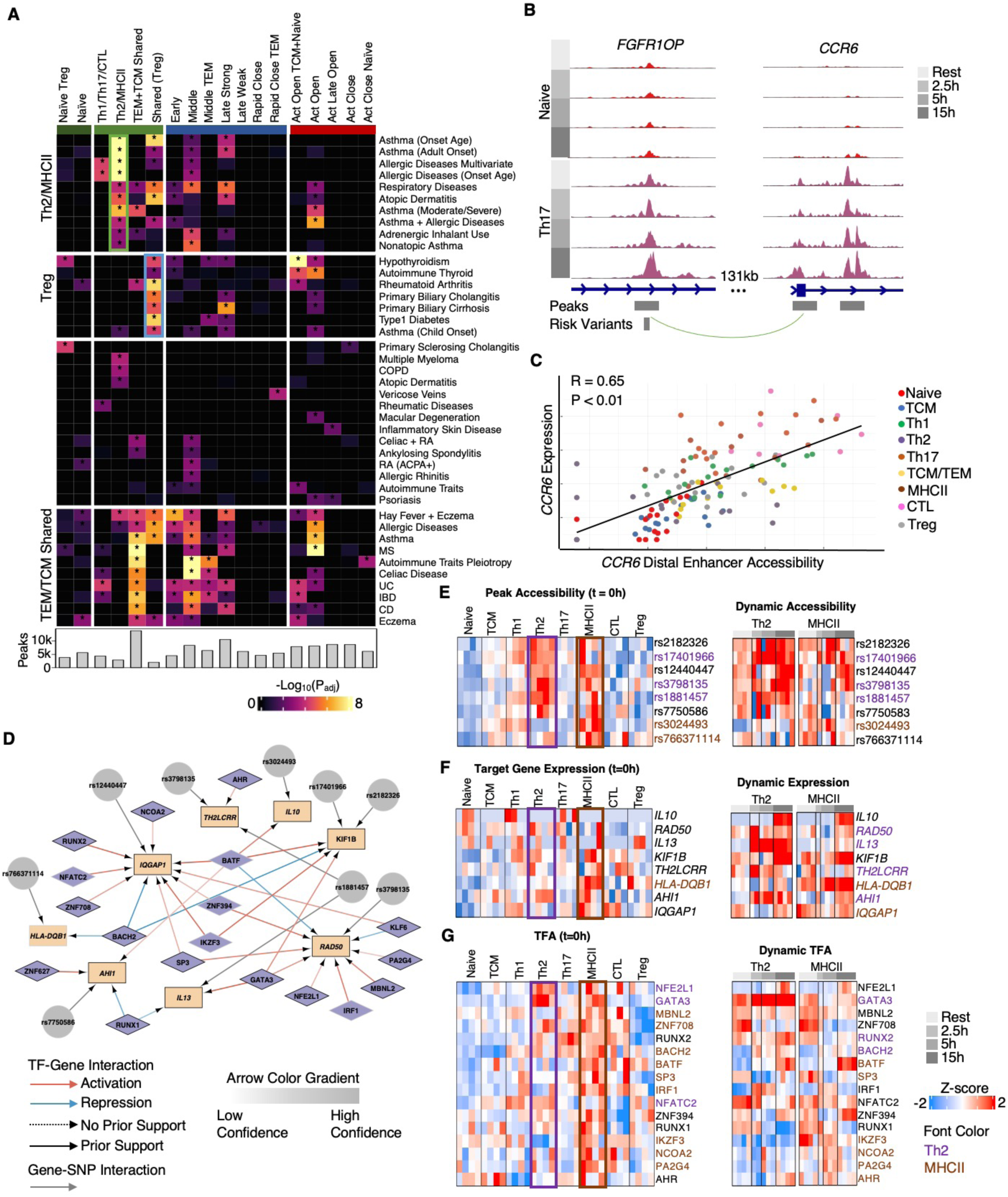
Integrated GWAS-GRN analyses nominate subset- and context-specific disease regulatory mechanisms in CD4+ T cells. **(A)** Significant colocalizations of GWAS phenotype genetic risk variant sets and accessible chromatin regions (resting memory- and naïve-associated, RR and those with naïve-memory shared activation dynamics); *P_adj_ < 0.1 (**Methods**). The clustered phenotypes are annotated based on the peak sets with the most enrichments (e.g., “Treg”). Green and blue boxes indicate risk variant sets selected for downstream GRN analysis. The size of each peak set is indicated. **(B)** The PSC risk variant (rs12529876), within the *FGFR1OP* locus, is linked to the *CCR6* promoter via track-looping 131kb away. The variant overlaps a RR peak (“middle TEM” dynamics) that is also a resting memory-associated peak (“TEM+TCM shared” cluster). **(C)** The accessibility of the distal *CCR6* enhancer in (B) correlates with *CCR6* expression (R = 0.65, P < 0.01). **(D)** GRNs centered on target genes linked (via trac-loop, micro-C and promoter proximity) to genetic risk variants in the “Th2/MHCII cluster” (green box in A). Only genes detected in >5% of Th2 or MHCII cells were included. TFs were included if they exhibited Th2 or MHCII-specific activity at rest or post-activation. Outlined nodes represent Th2 of MHCII RR genes. (**E**) Peak accessibility at risk variants, (**F**) target gene expression, and (**G**) TF activity of regulators. For panels E-G, the left heatmap represents the resting state (t=0h), while the right heatmap shows activated dynamics. Font colors indicate Th2-specific (purple) and MHCII-specific (blue) peak accessibility (E), gene expression (F) and TFA (G).

Clustering of disease-peak set enrichments yielded four phenotype groups **(Fig. 7A).** The “Th2/MHCII disease” cluster had the strongest enrichments in peaks defined by high accessibility in resting Th2 and MHCII cells. Nearly all diseases in this group were atopic, allergic or respiratory, underscoring the well-established role of type 2 immunity in these conditions. The “Treg disease” cluster, which included the autoimmune and inflammatory conditions (e.g., hypothyroidism, rheumatoid arthritis, type 1 diabetes), were most enriched in resting memory-associated peaks that were also accessible in Treg (other resting memory-associated peak sets were elevated in other memory populations but not Treg). Primary biliary cholangitis (**PBC**) was in the Treg-driven disease group; in support, PBC has been associated with an altered Th17/Treg balance(Kudira et al., 2024; R. Wang et al., 2024). A third cluster involved less significant enrichments, lacked coherent patterns and was not analyzed further. The fourth cluster, “TEM/TCM shared diseases”, encompassed both allergic diseases, such as hay fever and asthma, and inflammatory diseases, including ulcerative colitis, Crohn’s disease and celiac disease. Although diseases in this cluster most significantly overlapped resting memory “TEM+TCM shared” peak sets, they were also enriched for RR peak sets (“middle open”, “late open strong”) suggesting that RR dynamics might be impacted in these diseases. A subset also overlapped peaks with naive-memory shared opening dynamics (“act open“).

To explore potential CD4+ T subset-specific gene regulatory mechanisms for these diseases, we linked risk variants co-localized with accessible chromatin to genes, using promoter-enhancer interactions from the Trac-loop and Micro-C data to recover distal interactions and linear proximity (+/-1kb of TSS) for gene-proximal interactions. Using this method, 60% of overlapped risk loci were linked to putative target genes. Within an intron of *FGFR1OP,* the PBC-associated risk variant rs12529876 linked to the *CCR6* promoter 131kb downstream (**Fig 7B**). In patients with PBC, the frequency of CCR6^+^ immune cells in the hepatic portal tracts strongly correlated with the severity of inflammation (M. Zhang et al., 2024). Our analysis positions rs125892876 within a memory-associated accessible chromatin region, exhibiting elevated accessibility in resting memory populations (belongs to the “TEM+TCM Shared” cluster) and RR dynamics (“Late open strong”). In support that this putative enhancer drives *CCR6* expression, its accessibility correlates with *CCR6* expression across CD4+ T populations over the activation time course (**ρ** = 0.65, P < 0.01, **Fig. 7C**). Chemokine receptor CCR6 and its ligand CCL20 play important roles in several other inflammatory diseases, including inflammatory bowel disease, rheumatoid arthritis and psoriatic arthritis(Roghani et al., 2023; Shi et al., 2021; Varona et al., 2003). CCR6 may drive these diseases by enhancing Th17 migration to tissue lesions(C. Wang et al., 2009). In our data, both *CCR6* expression and the accessibility of its putative enhancer are highest in Th17 and the Th17-polarized CTL populations.

After linking risk loci to genes, we used the GRN to predict underlying regulatory mechanisms, focusing on risk variants for the first disease cluster (asthma, atopic respiratory) that co-localized with resting memory Th2/MHCII-defining accessible chromatin regions (**Fig. 7A** green box, **7D-E**). Of the thirteen risk-variant-linked genes, eight displayed Th2- or MHCII-specific expression at rest or upon activation (**Fig. 7F**), and we queried the GRN to identify TF regulators with elevated activity in the Th2 and MHCII populations at rest (**Fig. 7G**). The genes elevated in resting MHCII and/or Th2 populations *(KIF1B*, *TH2LCRR*, *HLA-DQB1, AHI1, IQGAP1)* also exhibited MHCII and/or Th2-specific dynamics upon activation, suggesting potential enhancer activity for the risk-variant linked putative regulatory regions. *RAD50*, *IL10* and *IL13* were not elevated in resting Th2 or MHCII but exhibited Th2-specific RR dynamics, suggesting a role for the risk-variant linked enhancers in Th2 poising of these genes. The GRN predicted known and novel regulators of these risk-variant-linked genes. For example, GATA3 is a well-known *IL13* regulator, while RUNX1 is novel. The novel interaction between KLF6 and the Th2 cytokine locus is supported by a ChIP-seq peak at *RAD50* within the locus (**Fig. 6I**). All regulators of MHCII-cell-associated risk genes are novel, given the novelty of this population (relative to well-studied Th2).

As a second example, we explored risk variant mechanisms for the second disease cluster (autoimmune and inflammatory conditions) enriched in resting memory Treg-defining accessible chromatin regions (**Fig. 7A** blue box, **S7**). Of the seven risk-linked genes, six displayed Treg-specific expression (**Fig. S7C**). *IL2RA, CTLA4*, *ZNF292* and *ZC3H12D* exhibited both Treg-specific expression at rest and upon activation, suggesting Treg-specific activity for the linked risk variant-containing regulatory elements. FOXP3, PRDM1, XBP1 and ATF4 were predicted to maintain baseline expression and RR poising of these risk-associated Treg genes (**Fig. S7D**).

Our analysis of GWAS variants connected disease-associated polymorphisms to diverse and dynamic accessible chromatin regions defining T cell states. Within CD4+ T cell populations and contexts, the GRN enabled identification of gene regulatory mechanisms that potentially contribute to genetic risk and pathogenesis of inflammatory and autoimmune diseases.

## Discussion

Memory CD4+ T cells protect against repeat infections via rapid cytokine production and clonal expansion upon antigen re-exposure. While many molecular components (receptors, TFs and epigenetic features) have been reported, the regulatory networks governing memory responses and rapid recall are poorly understood. Bulk ‘omics studies highlighted the role of the epigenome in regulating memory CD4+ T cell responses(Avni et al., 2002; Barski et al., 2017; Cuddapah et al., 2010; Fields et al., 2002) but were unable to resolve regulatory elements active in heterogeneous subsets. In this study, sn-multiome-seq enabled high-resolution, multi-modal analysis of CD4+ T cell populations, especially the TEM compartment, which was resolved into five populations: Th1, Th17, Th2, MHCII and CTL. Paired transcriptome-accessibility data were critical to our resolution of resting Th subsets, as well as heterogeneity in TCM and naïve cells. In our previous studies(Barski et al., 2017; Yukawa et al., 2020), memory T cell populations possessed positive chromatin marks proximal to rapid recall genes (including subset-specific genes, *IFNG*, *IL4*, *IL17A)*. With multiome-seq resolution, we confirmed that these resting chromatin accessibility regions were indeed subset-specific. The integration of chromatin accessibility and gene expression allowed us to reconstruct subset-specific signatures with higher precision than previously possible.

Multiome-powered GRN inference revealed novel regulatory circuits governing rapid recall responses in memory CD4+ T cells. In complex mammalian settings, integration of TF binding site (**TFBS**) predictions from chromatin accessibility improves GRN inference from gene expression data(Duren et al., 2017; Miraldi et al., 2019). Here, in a first-time application to GRN inference, we utilized maxATAC, cutting-edge deep neural networks(Cazares et al., 2023) to generate genome-wide “*in silico* ChIP-seq”, from snATAC-seq, for hundreds of TFs in our sc-resolved CD4+ T populations – a scale and resolution infeasible with existing TFBS measurement technologies. As a resource, the context-dependent, subset-resolved *in silico* ChIP-seq predictions are released as a UCSC Genome Browser Track Hub (Perez et al., 2025), for exploration alongside the corresponding chromatin accessibility measurements. These TFBS predictions, along with experimentally measured promoter-enhancer interactions(B. Lai et al., 2018; Oguchi et al., 2024), enhanced genome-scale GRN inference from the multiome-seq data. From the GRN, we predicted distinct and overlapping combinations of TFs orchestrating transcriptional and chromatin accessibility activation response dynamics (e.g., early versus late, subset-specific, memory-associated rapid recall and naïve-memory shared).

CD4+ T cells underwent extensive transcriptional and chromatin reprogramming following TCR activation, particularly during the early-to-middle activation phase (2.5-5h). In this window, cells upregulated effector molecules and their protein synthesis machinery. GRN analysis predicted known (MYC, XBP1, IRF4) and novel (MAX, NRF1) regulators of these processes. The wave of regulators between 5 and 15 hours (e.g., VDR, NFE2L1, PLAGL1, ATF5) controlled a distinct set of biological pathways, facilitating cellular homeostasis through induction of ER stress response and proliferation programs.

Our analysis implicated TFs (PRDM1, RUNX2, KLF6, MAF, SMAD3, NRN2C2, RORA, RORC) in memory maintenance at rest. As an example, we predict that SMAD3 promotes memory cell survival through *IL7Ra* regulation but suppresses RR pathways prior to activation, suggesting a checkpoint role preventing premature T cell responses. These memory-encoding factors orchestrate diverse programs targeting survival, cell-cycle and signal transduction genes, including enhanced expression of costimulatory molecules other than *CD28*. Upregulation of *CD58* and its binding partner *CD2* might contribute to stronger activation responses and RR kinetics of memory T cells *in vivo*. Excepting RUNX2 and SMAD3, these TFs maintain high activity post-activation, contributing to enhanced cytokine production in addition to memory maintenance.

We sought support for these and other GRN predictions through evaluation of external datasets. Our GRN was constructed from multiome-seq data derived from four independent donors. Thus, evaluation and refinement of our predictions in a larger cohort, scRNA-seq of similarly aCD3/CD28-activated CD4+ T cells (n = 119 individuals) was invaluable (Soskic et al., 2022). For example, most but not all of our GRN-predicted memory-core TFs were supported by the larger study, leading to refinement of our model (**Fig. 5F-G**). The larger study was also used to confirm TF activity dynamics (**Fig. S3H**, **S4B**), the novel TCM populations (**Fig. S4E-F,I**) and their underlying regulation (**Fig. 3G-H**). Equally important, the larger cohort enabled examination of biological variables (sex and age). We identified significant age-dependence of TF activities within the naïve but not memory cell compartments (**Fig. S4D**). Memory factors associated with rapid recall potential, KLF6 and PRDM1, were age-increased within the naïve compartment, but these correlations, although significant, are minor sources of variation relative to activation- and cell-type-dependent differences (e.g., as shown in **Fig. S5F**), suggesting that the definition of our memory core is likely to be robust across age groups as well as sex. We also established support for individual regulatory interactions through generation of ChIP-seq data for the novel memory-associated factor KLF6 and curation of TF ChIP-seq and perturbation data for MAF and PRDM1 (**Fig. 6H,I**).

We also explored whether the GRN predictions could be extrapolated to more physiological, antigen-specific CD4+ T cell activation contexts. For this purpose, we utilized an scRNA-seq study involving peanut-stimulation of CD4+ T cells from peanut allergic individuals undergoing oral immunotherapy (Monian et al., 2022). For our memory-associated core of five TFs (MAF, PRDM1, KLF6, RUNX2, SMAD3), there was good qualitative agreement between TF activities from the antigen-specific data and our GRN simulations (**Fig. 6J**). Extension of the analysis to a larger, unbiased set of activation-dependent TFs again identified many TFs with qualitatively similar behavior (70%) but also TFs with divergent TFA predictions, particularly those with high activity in peanut-activated Treg (**Fig. S6**). These differences are likely due to expected differences in molecular biology of allergen-specific T cells and their activation, as well as limitations of the GRN model simulations. Given the diversity of T-helper polarization environments induced by different antigens, evaluation and refinement of the GRN with data from physiological settings is a critical future direction. Comparative analyses provide a framework to identify regulatory mechanisms distinguishing antigen-specific activation from general T-cell stimulation. Our GRN thus serves as a base model to dissect distinct modes of T-cell activation and explore how context-dependent signaling shapes effector function.

Comparison of GRNs across *in vivo* and *in vitro* contexts also will be a valuable means to identify limitations of the aCD3/CD28 T cell activation model and an opportunity to iteratively design better *in vitro* human models (e.g., via identification of key regulatory interactions driving cell behavior *in vivo* but missing *in vitro*). Indeed, many genomics assays (including the TF binding and promoter-enhancer interaction measurements used here) require enrichment strategies and a large number of cells, limiting their direct application to complex physiological settings and creating a need for reliable, well-characterized and reproducible *in vitro* and *in silico* human models(Forlin et al., 2023; Hartung & Kleinstreuer, 2025). Thus, our GRN also serves as a base model to aid in the development of higher-fidelity models of human CD4+ T cell activation.

In future work, the GRN may enable identification of antigen-specific activated memory and naive CD4+ T cells from *in vivo* sc-genomics, including spatial transcriptomics, studies (i.e., where cell enrichment strategies may not be feasible and/or the number of transcripts detected may be limited). In the *Monian et al.* peanut-activation study, cells were flow-sorted to enrich for peanut-specific CD4+ T cells from blood. We highlight that our GRN was more successful predicting TF activities distinguishing peanut-activated memory cells (CD154^+^, CD137^+^) from resting (DN) cells than subtler differences between the activated TEM and Treg populations (**Fig. S6**), suggesting that GRN-based estimation of TF activities could be developed further to identify individual antigen-specific activated T cells from *in vivo* data.

Finally, the GRN may have utility deciphering impacts of human genetic variation on T cell behavior. Encouraged by the benchmarking, we explored the GRN for potential regulatory mechanisms underpinning noncoding genetic risk in CD4+ T cells. The resting memory core TFs may mediate genetic risk for immune-mediated disease, as accessible chromatin maps defining the resting memory subsets (Th2/MHCII, Treg, pan-TEM/TCM) were most frequently and significantly associated with genetic risk variants for immune-mediated diseases (**Fig. 7A**). Focusing on atopic and asthma variant colocalizations with the resting Th2/MHCII-associated chromatin, we identified regulators, including RUNX2 and KLF6, of the risk-variant target genes (**Fig. 7D**).

In conclusion, we describe a genome-scale GRN governing rapid recall and early activation timepoints in human naïve and memory T cell subsets. This sn-multiome-inferred model provides a foundation for future studies aimed at deciphering the transcriptional networks governing T cell responses in health and disease. In this manuscript, we highlight but a subset of regulatory mechanisms from the GRN and rich, underlying data. To maximize the resource, the GRNs and sn-multiome data are interactively available, along with open-source codebases to reproduce results here-in, from: https://github.com/MiraldiLab/RapidRecall.

## Methods

### Cell processing

Five leukocyte depletion filters containing cells from de-identified male donors were obtained from the University of Cincinnati Hoxworth Blood Center, Cincinnati, Ohio. Filters were back-flushed with PBS, and PBMCs were prepared by Ficoll centrifugation. Human CD4+ T cells were isolated from PBMCs using CD4 isolation magnetic beads (STEM CELL #19052). To increase the fraction of TEM cells, naive and TCM cells were partially depleted using anti-CD27-biotin coated streptavidin magnetic beads (Invitrogen Dynabeads Biotin Binder 11047; CD27 antibody Biolegend 302804). Cells were cryopreserved in 50% CTS OpTmizer medium (A1048501), 40% human serum (Sigma H4522), 10% DMSO (ThermoFisher) and stored in liquid nitrogen. Cells were prepared for 16 multiome-seq experiments (4 donors x 4 time points). Thawed, purified cells were plated in CTS OpTmizer with 1% Penicillin-Streptomycin (Gibco 15140122) and incubated at 37°C for the activation duration time: 15h/5h/2.5h. Activation was performed using Dynabeads Human T-Activator CD3/CD28 (Gibco 11131D). Resting cells were incubated in the same medium and temperature for the same time as activated, but without activation beads added. To prevent cell-free DNA contamination originating from dead cells, DNase I (1:100) was added directly into the medium (NEB M0303S) during the last 30min of incubation. After incubation, cells were washed with PBS + 0.1% BSA twice. For each multiome experiment, cells from the spike-in control donor (resting timepoint) were mixed with cells from the experimental donor at the time point of interest at a ratio of 8% (ctrl) to 92% (each test donor), immediately preceding the nuclei prep. Nuclei were prepared according to the modified 10X Genomics protocol: 100ul of cold lysis buffer (10mM Tris-HCl (pH 7.4), 10mM NaCl, 3mM MgCl2, 1% BSA, 0.025% Tween-20, 0.025% Nonidet P40, 0.00125% Digitonin, 1mM DTT, 1 U/µl RNase inhibitor (Roche)) was added to cell pellets. Samples were incubated for 40s on ice, washed with 1mL of cold wash buffer (10mM Tris-HCl (pH 7.4), 10mM NaCl, 3mM MgCl_2_, 1% BSA, 0.025% Tween-20, 1mM DTT, 1 U/µl RNase inhibitor (Roche)) and filtered with Flowmi cell strainer (Bel-Art H13680-0040).

### scRNA and scATAC libraries preparation

The Chromium Next GEM Single Cell Multiome ATAC + Gene Expression assay was performed according to the manufacturer’s instructions (10x Genomics) with the Chromium Next GEM Single Cell Multiome ATAC + Gene Expression Reagent Bundle (PN-1000283) kit. Briefly, nuclei suspended in the multiome buffer were combined with a transposition mix and incubated for 1h. The transposition reaction was combined with the kit’s master mix and loaded together with partitioning oil and gel beads into the Chromium Next GEM Chip J Single Cell Kit (PN-1000234) to generate a gel bead-in-emulsion (**GEM**). The GEMs were incubated to produce 10x-barcoded DNA from the transposed fragments (for ATAC) and 10x-barcoded, full-length cDNA from poly-adenylated mRNA. Next, the GEMs were broken, the barcoded DNA and cDNA were purified, and the purified products were subjected to a pre-amplification PCR step. The pre-amplified material served as starting material for ATAC and RNA library construction per 10x protocol. ATAC libraries and GEX libraries were pooled separately. The pools were sequenced on the NovaSeq 6000 sequencer with SP or S1 flow cells, using the following sequencing parameters: R1: 50 cycles, i7: 8 cycles, i5: 24 cycles, R2: 49 cycles (for ATAC) or R1: 28 cycles, i7: 10 cycles, i5: 10 cycles, R2: 90 cycles (for RNA).

### Whole genome sequencing and variant calling

DNA was isolated using a PureLink Genomic DNA Kit (Thermo Fisher). Whole-genome sequencing was performed using DNBseq next-generation sequencing technology. Libraries were sequenced on an Illumina NovaSeq to generate 150-base paired-end reads. Sequencing reads were aligned, and variants were identified using the Genome Analysis Toolkit (**GATK**) Unified Genotyper following GATK Best Practices(DePristo et al., 2011; McKenna et al., 2010; Van der Auwera et al., 2013).

### scRNA-seq and scATAC-seq quality control

Initial alignment, filtering and UMI counting were performed using cellranger-arc-2.0.2 and the 10x genome GRCh38 reference genome within the Scientific Data Analysis Platform (https://scidap.com)(Kotliar et al., 2025). Then, using Seurat v5.1.0, the following quality thresholds were used to select quality cells: detected genes > 400, transcript unique molecular identifiers (**UMI**) > 750, percent mitochondrial transcripts < 20%, ATAC fragments > 2000, fragments in blacklist < 5%, nucleosome signal < 2.5, TSS enrichment > 2(Hao et al., 2021). Initial clustering revealed small populations of non-CD4+ T cells, including Natural Killer, CD8+ T and myeloid cells based on either *CD8A* or lack of *CD3E* expression. These cells were removed from the analysis. We then used several criteria to filter likely doublet nuclei. We removed nuclei (1) predicted to be doublets based on both RNA (scDblFinder v1.15.2) and ATAC (ArchR v1.0.2) modalities and (2) those with unexpectedly large RNA (>10k UMI) and ATAC (>50k fragment) libraries(Germain et al., 2021; Granja et al., 2021). For scDblFinder, a merged SingleCellExperiment object was used for input (additional params: Cluster = TRUE, samples = “Dataset“). For ArchR, the 16 individual fragment files were used as input, as well as each samples expected number of doublets. The expected number of doublets was calculated by multiplying the average doublet rate reported by the Cell Ranger User Guide(*CellRanger Tutorial*, n.d.) by the number of cells in the sample. Across the 16 multiome-seq experiments, our QC pipeline yielded 65,862 quality, CD4+ T cell nuclei.

### Sample integration and clustering

Prior to sample integration, each sample’s Cell Ranger fragment file was modified to include both transposase cut sites(Cazares et al., 2023; Wayman et al., 2023). Cut sites were filtered if they overlapped ENCODE GRCh38 blacklist regions. Peak calling was performed by MACS2 V2.1.4(Gaspar, 2018; Y. Zhang et al., 2008) on initial clustering of each Seurat dataset individually (MACS2 parameters: --shift -25 --extsize 50 --tsize 50 --pvalue 1E-8 --keep-dup all)(Wayman et al., 2023). A reference peak set was generated by taking the union of peaks called across all samples and merging overlapping peaks. A (peaks x cells) counts matrix was then generated using *FeatureMatrix* from Signac v1.13.0 (Stuart et al., 2021). This counts matrix replaced the original counts data generated from Cell Ranger.

Next, the Seurat v5.1.0 pipeline was applied to the RNA and ATAC data in parallel. For RNA, the data was log-normalized, centered and scaled by z-scoring, and then dimensionality reduced to the first 30 PCs. For ATAC, data was first normalized using run term frequency inverse document frequency (the *RunTFIDF* function) followed by latent semantic indexing (**LSI**) dimension reduction (the *RunSVD* function) using components 2-30. Component 1 was removed due to high correlation with library size (fragments per nucleus). Peaks with ≥1 cut site in 20 or more cells were included in the dimension reduction. After dimensional reductions for both RNA and ATAC were calculated, each data modality was then separately batch-corrected using Harmony v3.8(Korsunsky et al., 2019). The batch variable provided to Harmony was dataset (1-16), with either the PCA or LSI reduction as input. The output of Harmony is a batch-corrected data dimension reduction (harmony_rna, harmony_atac). The two data modalities were integrated using Seurat’s weighted nearest neighbor (**WNN**)(Hao et al., 2021) method using the Harmony dimension reductions as input. Cells were clustered using the smart local moving (**SLM**) algorithm. Other clustering algorithms (Louvain and Leiden) gave comparable results. After clustering, another round of peak calling was performed with MACS2 (see above parameters), with cells aggregated per cluster-timepoint. These peak sets were merged (union) to generate a penultimate reference peak set. The above ATAC-seq steps (normalizing, dimension reduction, harmony integration) and combination with transcriptome (wnn, clustering) were repeated to achieve the final clustering solution: 15 clusters and 65,862 cells. Cluster identities were determined based on gene markers and accessibility near effector genes and genome-scale analyses, as described in **Results** (**Fig. 1C-E**, **S2**). A cluster of “stressed” cells (3.1% of total) bore markers of cell stress (*TXNIP*, *PDCD4*), lacked activation markers (*CD44, IRF4*) and was excluded from downstream analyses.

### Identification of “Donor0” spike-in control cells

To distinguish the spike-in control (“Donor0”) from experimental donor cells in each multiome-seq experiment, we used genetic variant calls, available from our WGS of the spike-in control (Donor0) or inferred from the RNA and ATAC multiome modalities for the four experimental donors, using the standard GATK pipeline (v4.2.4.1)(Betschart et al., 2022). First, we identified high-confidence experimental donor cells from each sample using Souporcell v2.5(Heaton et al., 2020). Souporcell was run independently for the RNA and ATAC data, using Cell Ranger BAM files as input. To improve Souporcell speed and performance, we provided a single nucleotide variant (**SNV**) reference of common variants occurring in at least 5% of individuals from the 1000 Genomes Project(1000 Genomes Project Consortium et al., 2010); additional Souporcell parameters included: --skip_remap True, --ignore True, -k 2, --no_umi True (ATAC only). Second, to further improve genotype-based mapping of cells to experimental donors (lacking WGS), donor-cell predictions supported by both RNA and ATAC analyses were used to infer genotypes for the experimental donors, serving as valuable input to a second round of Souporcell analysis. For variant calling in these experimental donors, the Cell Ranger BAM files from each sample were filtered based on predicted experimental cell barcodes (supported by RNA and ATAC) and variants were called using the standard Freebayes v1.3.7 pipeline(Garrison & Marth, 2012) applied to the RNA sequencing data. Finally, another round of Souporcell was performed on each sample, this time using the full gene expression BAM file and each donor’s predicted VCF files as input.

### Pseudobulk batch correction, data normalization and differential analyses

We aggregated gene expression or accessibility signal from individual cells to generate pseudobulk profiles per CD4+ T cell subset, time point and donor. Importantly, the spike-in control donor cells provided a pseudobulk sample condition (“Donor0”, resting cells) that remained constant across all 16 multiome-seq experiments (donor-timepoints). This batch-control design is required to distinguish biological from technical variation across experiments, as standard batch correction methods (ComBat, DESeq2, EdgeR)(Johnson et al., 2007; Love et al., 2014; Y. Zhang et al., 2020) require at least one common control sample across experiments. However, the use of spike-in donor control cells is not common practice in sc-genomics study designs. Inclusion of a spike-in donor (8% of nuclei per experiment) (1) slightly reduces sequencing data available for the experimental donor-condition of interest (i.e., incurs expense) and (2) the processing of the spike-in cells introduces additional steps and therefore risk to sensitive nuclei prep protocols, which may not be feasible or acceptable for, e.g., challenging tissue samples or rare patient samples.

We therefore evaluated the impact of our spike-in study design on inference quality, both visually and quantitatively. As described above, single-cell Harmony batch correction was critical to resolving nuclei across experiments into cell populations (**Fig. S1A-C**). The pseudobulk gene expression counts resulting from Harmony batch-corrected cell integration and WNN clusters served as input to three pseudobulk batch-correction approaches: (1) no pseudobulk batch correction, (2) spike-in informed batch correction, where each of the 16 multiome-seq experiments was treated as a batch with a common pseudobulk (spike-in) sample, (3) batch correction ignoring the spike-in control pseudobulk (for we supplied an incorrect design in which the four timepoint experiments per donor were treated as a single batch (“batch = donor”), resulting in 4 batches that preserved timepoint information). Option (3) is commonplace in the absence of spike-in control cells, but problematic for several reasons: (a) Biological variability due to donor differences is modeled as technical and subtracted off as a batch effect. (b) Despite targeting similar numbers of nuclei for input to the 10x instrument, the number of quality nuclei and their per-nucleus library sizes are often variable across experiments. Lumping four experiments into a single batch, based on donor, makes it difficult to control for this important technical covariate (among others). For batch correction, we used Combat-Seq from sva v3.46.0 (Leek et al., 2012), using “option 2” spike-in informed parameters (batch=dataset, group=celltype-timepoint) or “option 3” parameters (batch=donor, group=celltype-timepoint).

For data normalization and differential analysis, we used DESeq2 (v.1.38.3) and the variance stabilizing transformation (**VST**). We also leveraged the single-cell nature of our data for differential gene expression analysis, filtering genes that met our DESeq2 statistical cutoffs (FDR=10%, log_2_|FC|>0.58) but were detected in fewer than 5% of nuclei within one condition of a comparison (2.5h activation versus resting in Th2 cells). As previously(Wayman et al., 2023), this additional sc-based differential criterion improved biological signal, removing low-quality differential genes supported by pseudobulk analysis alone. For differential analysis of the accessible chromatin data, we performed no filtering based on % nuclei with peak signal, because peak signal per nucleus is limited by DNA copy number and extreme signal from a small number of cells cannot occur. This rationale was supported by visual inspection of ATAC signal quantification (**Fig. 4A, 5A, S5A**). For differential analysis of ATAC signal, FDR=10%, log_2_|FC|>0.58 were also applied.

To visually compare the three pseudobulk batch-correction strategies, we identified the top 1000 most batch-dependent dynamic genes resulting from the “no batch correction” strategy for visualization. First, from the “no batch correction” analysis, dynamic genes were identified as those differential for at least one activation timepoint relative to t=0, for at least one cell type. Next, we identified those dynamic genes with the greatest batch effects, using the following strategy: (1) We calculated the coefficient of variation (**CV**) for each gene at each celltype_timepoint across the four donors, using VST counts from the “no batch correction” strategy. (2) For each gene, the CVs were averaged, and the 1000 genes with the highest average CV were visualized in the heatmap (**Fig. S1D-F**). As expected, performing no pseudobulk batch correction resulted in the largest batch effects. We quantitatively assessed the three pseudobulk batch-correction strategies by comparing the number of differential genes resulting from each strategy. In addition, we considered that the incorrect “batch = donor” assumption is only likely for experiments lacking spike-in control cells, and, in that case, a “batch = donor” design would benefit from additional experimental donor cells (i.e., a slight signal gain, as the experimental donors would contribute 100% (as opposed to 92%) of cells in the experiment). For this reason, we added a simulation condition to our comparisons, randomly replacing spike-in donor cells with experimental donor cells for each experiment. (Similar to bootstrapping, our up-sampling simulation underestimates gene variance, providing a challenging comparator for the spike-in strategy.) For all comparisons, the spike-in batch control strategy resulted in the most differentially expressed genes. The relative increase in DEGs was dependent on the strength of the biological signal: While the spike-in strategy increased activation-dependent DEGs by 8-12%, depending on cell population timepoint (**Fig. S1G**), the spike-in strategy increased DEGs between difficult-to-distinguish memory populations at rest by 20-30% (**Fig. S1H**). Not only do these results demonstrate the benefit of the spike-in design in our study, but they also provide support for this batch-control approach in future sc-genomics studies.

For the scATAC-seq data, the same processing steps were followed, with the exception that we did not run ComBat-seq to control for batch effects, as visual inspection of the data (analogously to **Fig. S1D** analysis) did not identify gross batch effects.

Pseudobulk samples that were too sparse (library size <150k for RNA, <400k for ATAC) were excluded from downstream analyses, with the following exception: at the 2.5h timepoint, no donor met the RNA or ATAC cutoff for the Th2 and Th17 populations. Given the importance of these populations at this time point, we took the following approach: Following batch correction with ComBat-seq (RNA) and proceeding DESeq2 normalization, RNA counts were combined for donors “1 and 2” and donors “3 and 4”, resulting in two pseudobulk gene expression and chromatin accessibility estimates at 2.5h for Th17 and Th2 cells.

For visualization of pseudobulk chromatin accessibility tracks (**Fig. 1E**), data were library size normalized following the method used in Signac(Stuart et al., 2021).

### Dendrograms

Raw pseudobulk RNA and ATAC counts matrices were downsampled to 150,000 reads and 400,000 fragments, respectively, to account for differences in library sizes across populations. Following the batch correction and normalization strategies describe above, downsampled counts matrices were batch-corrected (RNA only) and VST-normalized prior to z-scoring (both). Features with a coefficient of variation of < 0.01 for expression and < 0.001 for accessibility were filtered out, resulting 12,225 genes and 23,136 ATAC peaks. Hierarchical clustering was performed using *ComplexHeatmap* (v 2.14.0)(Gu et al., 2016) with Ward’s method and Euclidian distance.

### Identification of rapid-recall genes and accessible chromatin regions

Rapid recall genes or peaks were defined as features with faster, more extreme activation dynamics (up or down) in memory populations than in naive cells. Therefore, a feature is rapid recall if:

1. It is dynamic in at least one memory population (differential at 2.5h, 5h or 15h relative to t=0h) AND
2. at the above dynamic timepoint(s) in the corresponding memory population(s), the feature is differential relative to naïve cells.
3. The sign of the log_2_(FC) estimates for (1) and (2) are the same (thereby ensuring that at least one memory population has faster, more extreme dynamics than naïve).

Differential analysis criteria are as described above.

### GSEA

We conducted gene set enrichment analysis (**GSEA**) using the Gene Ontology (**GO**) Biological Processes pathways obtained from the Molecular Signatures Database (**mSigDB**, accessed April 8, 2024)(Subramanian et al., 2005). Statistical significance was assessed using Fisher’s exact test, with the analysis restricted to pathways containing a minimum of 5 genes. The background gene set comprised all genes present in the above pathways. Pathway enrichments were considered statistically significant at a (**FDR**) threshold of 10%. Multiple testing correction was applied using the Benjamini-Hochberg procedure.

### Motif scanning and enrichment

We performed motif scanning using CIS-BP motifs (v2.0.0)(Weirauch et al., 2014) on all peaks in our reference set (described above) using FIMO from Meme Suite version 5.4.1 (raw P-value threshold = 0.00001, first-order Markov model background)(Grant et al., 2011). Enrichment of TF binding sites in peak sets compared to background (all peaks) was calculated using a Fisher exact test (FDR = 10%) and Benjamini-Hochberg procedure to correct for multiple hypothesis testing.

### Cell cycle scoring

To score cell-cycle phase for each cell, we downloaded genes associated with each cell phase from tinyatlas (update from 9/13/17)(Kirchner & HBC, 2018). We then used Seurat’s *CellCycleScoring* function to assign scores at the single-cell level, using the phase with the highest score to classify cells.

### GRN model construction

#### Selection of Target Genes for GRN

We selected 15,169 target genes for GRN inference, based on differential expression (FDR=10%, log_2_|FC|>0.58) within CD4+ T cell subsets across timepoints, as well as across subsets at a particular timepoint.

#### Selection of Potential Regulators

We intersected a curated list of 1639 human TFs(Lambert et al., 2018), with our list of target genes (above), yielding an initial set of 761 regulators. *RORC* did not meet our definition of a target gene but was added as a potential regulator, given its known importance to CD4+ T cell biology(Ivanov et al., 2006). *CTCF* was a target gene, but we removed it from consideration as a potential regulator(Bell et al., 1999), given its mode of regulation as an insulator. Thus, we considered a total of 761 potential regulators for GRN inference.

#### maxATAC prediction of TFBS

We used maxATAC (version 1.0.5)(Cazares et al., 2023) to generate genome-wide *in silico* TF binding site (TFBS) predictions for human TFs that met our potential regulator criteria and had maxATAC models available. To ensure biological relevance, we only included maxATAC predictions for TFs that were expressed in at least 5% of cells within the cell type of interest. This expression filtering resulted in TF-specific predictions for each of the fourteen CD4+ T cell populations, with the final set comprising 110 TFs: ARID3A, ARNT, ATF1, ATF2, ATF3, ATF7, BACH1, BHLHE40, CEBPB, CREB1, CREM, CUX1, E4F1, EGR1, ELF1, ELF4, ELK1, ESRRA, ETS1, ETV6, FOS, FOSL1, FOSL2, FOXK2, FOXP1, GABPA, GATA3, GATAD2B, HSF1, IKZF1, IRF3, JUN, JUNB, JUND, KMT2A, LEF1, MAFF, MAFK, MAX, MAZ, MBD2, MEF2A, MNT, MXI1, MYB, MYC, NFATC3, NFE2L2, NFIA, NFIC, NFXL1, NFYA, NFYB, NKRF, NR2C1, NR2C2, NRF1, PBX3, PKNOX1, RELA, REST, RFX1, RFX5, RUNX1, SETDB1, SKIL, SMAD5, SP1, SREBF1, SREBF2, SRF, STAT5A, TBP, TCF7, TCF7L2, TCF12, USF2, YBX1, YY1, ZBED1, ZBTB7A, ZBTB11, ZBTB33, ZBTB40, ZEB1, ZFX, ZHX2, ZKSCAN1, ZNF24, ZNF143, ZNF207, ZNF217, ZNF274, ZNF282, ZNF384, ZNF407, ZNF569, ZNF592, ZSCAN29, ZZZ3. We generated maxATAC predictions for each of the fourteen CD4+ T cell populations in **Fig. 1B**, as they all met the recommended minimum library size (5M fragments) for maxATAC prediction from snATAC-seq data. We followed default data processing of ATAC signal for input to the pre-trained maxATAC models. The total number of genome-wide “in silico TF ChIP-seq” tracks varied by population based on TF expression, with predictions generated only for TFs meeting the 5% expression threshold in each respective population. To arrive at a final set of TFBS, we prioritized precision over recall, thresholding maxATAC scores to achieve an expected recall of 10%, as reported per TF model (Cazares et al., 2023).

#### Association of putative regulatory regions (ATAC peaks) with target genes

Human CD4+ T cell Trac-loop data was downloaded from GSE87254(B. Lai et al., 2018). We combined unique Trac-loop regions from both resting and activated cells into one list. Genomic regions were lifted from hg19 to hg38 using UCSC’s Lift Genome Annotation tool with a minimum ratio of remapped bases of 0.95 (6 of 38812 regions could not be mapped to hg38) (Perez et al. 2025). To infer enhancer-promoter interactions in our data, we overlapped our reference ATAC peak set with Trac-loop regions directly connected to a gene promoter. Similarly, we downloaded the Micro-C data from DDBJ (ascension PRJDB13816 (Oguchi et al., 2024)) and combined all detected loops from both resting and activated cells. A loop was linked to a gene if either side of the loop overlapped a TSS for a gene. Then, if an ATAC-seq peak intersected either side of that loop, it was linked to that gene.

#### Prior network of TF-gene interactions inferred from ATAC data

A TF-gene interaction was included in the prior if a predicted binding site for that TF (derived from maxATAC or TF motif scanning, see above) overlapped an ATAC-seq peak (from our reference peaks set) that was within 5kb of the gene TSS or linked distally to the gene via Trac-loop or Micro-C (see above).

#### GRN inference

Gene expression is modeled as a multivariate linear combination of transcription factor activities (**TFAs**):

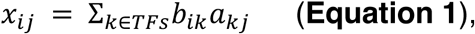

where *x_ij_* corresponds to the expression of gene *i* in pseudobulk sample biological replicate *j* (“resting Th1, Donor 2”), (*a_kj_* is the activity of TF *k* in condition *j*, and *b_ik_* describes the regulatory effect of TF *k* on gene *i* (Bonneau et al., 2006; Miraldi et al., 2019; Wayman et al., 2023)). While gene expression is directly measured in our multiome-seq study design, protein TF activities (*a_kj_*) are not. Based on previous GRN benchmarking in CD4+ T cells (bulk RNA-seq and ATAC-seq(Miraldi et al., 2019) and scRNA-seq and scATAC-seq(Wayman et al., 2023)) we used two complementary TFA estimation strategies:

(1) Prior-based TFA assumes that protein TFAs can be estimated based on the expression of a TF’s target genes, using partial prior knowledge of TF-gene interactions (as inferred from ATAC-seq and Trac-loop above). TFA is calculated from the normalized gene expression matrix *X* ∈ ℝ^|genes|×|samples|^ and the prior network matrix *P* ∈ ℝ^|genes|×|*TFs*|^ using the least-squares solution of the equation(Arrieta-Ortiz et al., 2015):

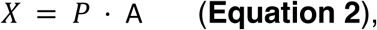

(2) TF mRNA serves as an estimate of protein TFA; this is a reasonable assumption for some TFs, whose major mode of regulation is transcriptional (as opposed to post-transcriptional).

For each TFA estimation strategy separately, TF-gene regulatory interactions, *b_ik_*, were identified via optimization of the following objective function(Miraldi et al., 2019):

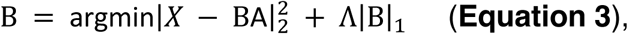

where *X* ∈ ℝ^|genes|×|samples|^ is the gene expression matrix, *A* ∈ ℝ^|TF^*^s^*^|×|samples|^ is the TF activity matrix, *B* ∈ ℝ^|genes|×|*TFs*|^ is the matrix of TF-gene interaction coefficients, and Λ ∈ ℝ^|genes|×|*TSs*|^ is the matrix of penalties associated with each interaction. The penalty matrix incorporates prior information via a decreased penalty (Λ*_ik_*) for interactions with prior support (Λ*_ik_* = 0.5 with prior support, Λ*_ik_* = 1 for no prior support). While the adaptive LASSO penalty encourages selection of TF-gene interactions supported by the prior, novel interactions, strongly supported by the gene expression model, can be inferred. In this way, the two sources of error in the prior (false positive and false negative interactions) can be corrected through the inference procedure. The ability to infer novel interactions is especially critical for TFs lacking a maxATAC or TF motif model (e.g., TET2) and therefore ATAC-derived prior information. The LASSO regularization parameter λ using StARS(H. Liu et al., 2010), with 220 subsamples and a subsample fraction of 0.05. Confidence scores for TF-gene interactions were defined as the sum of (1) subsamples in which the edge was nonzero and (2), to break ties between equally stable edges, the correlation between the TF’s activity and its predicted target gene’s TFA(Miraldi et al., 2019). We evaluated the GRN’s predictive performance (predictive R^2^ on left-out test gene expression data) as a function of model complexity (average number of regulators per gene)(Miraldi et al., 2019); this analysis identified an average of 10 regulators per target gene as optimal for GRNs derived using both prior-based TFA and TF mRNA. Confidences from these two networks were rank-combined using the maximum(Miraldi et al., 2019), then the top 151,140 interactions were selected and reranked and signed, using partial correlation between genes and their TF regulators to determine interaction sign.

### GRN analyses

#### Final transcription factor activity

estimates were calculated by substituting the signed, ranked confidences from the final GRN for the prior matrix in **Equation 3**. This procedure was applied to our dataset and also when evaluating TFA in the independent scRNA-seq studies(Monian et al., 2022; Soskic et al., 2022).

#### Functional annotation of TFs using the GRN (e.g., Fig S3D)

To determine whether a TF activated (or repressed) gene pathways, we tested for significant overlap between its positive (or negative) target genes and gene pathways, using GSEA (described above).

#### GRN subnetworks regulating proliferation genes in TCM. (e.g., Fig. 3D)

We first compared estimated transcription factor activity between TCM and TEM, with TEM activity represented as the average across Th1, Th2, and Th17 subsets. Transcription factors with an average z-score difference of at least 1.5 (higher in TCM) were retained. We then assessed these TFs for enrichment of their target genes in the Gene Ontology Cell Cycle pathway; since all TFs were enriched, none were excluded. Finally, we restricted the network to high-confidence regulatory interactions that were observed in a minimum of 200 out of 220 GRN construction subsamples (see above).

#### GRN subnetworks encoding TCM/TEM (e.g., Fig. 3H)

We selected TFs that were differentially active between TCM/TEM and TCM in the same direction at all 3 activation timepoints (FDR = 10%). Target genes for this network included genes upregulated in TCM/TEM compared to TCM in at least one activation timepoint.

#### GRN subnetworks encoding memory at rest (e.g., Fig. 5G)

TFs predicted to encode resting memory were prioritized based on several criteria:

1. The TF’s motif is enriched in a peak cluster with increased accessibility in memory versus naïve cells (e.g., **Fig. 5A** clusters Th1/Th17/CTL, Th2/MHCII)
2. The TF is differentially expressed between at least one memory population and naïve cells at rest
3. The TF has at least two predicted upregulated target genes (positive edge) that show differential expression between naïve cells and at least one memory population at rest, where the expression is higher in the memory population.
4. The TF had significantly higher estimated activity in TEM cells compared to naive in soskic et al.

Nine TFs met these criteria:PRDM1, RORC, RUNX2, NR2C2, KLF6, RORA, SMAD3, MAF. Target genes were upregulated in memory cells compared to naive at rest and involved in the biological pathways enriched in memory upregulated genes (**Fig. 5E**).

#### Disease GRN subnetworks

We constructed GRNs centered on the accessible chromatin peak set most significantly overlapping the “Th2/MHCII” or “Treg” phenotype variant sets (green and blue boxes in **Fig. 7A**), to arrive at Th2/MHCII (**Fig. 7D**) and Treg (**Fig. S7A**) disease GRNs. We first collected all risk variants with an overlapping peak from the peak-cluster of interest (Th2/MHCII or Shared Treg). These peaks were linked to genes using the method described in *Prior network of TF-gene interactions inferred from ATAC data*, with the exception that the proximity rule was based on a more stringent criterion +/- 1kb promoter (**Table S9**). Any SNP-linked target genes detected in < 5% cells of the relevant cell-type (Th2/MHCII or Treg) were excluded from the analysis. We then visualized the expression dynamics of the target genes as well as the activities of all predicted TF regulators from the GRN. Target genes and their TF regulators were retained in the GRN if their expression and TFA were elevated in the cell-type of interest at rest or their activation dynamics were specific to the cell-type of interest.

### Modeling chromatin accessibility as a function of TF activities

We built a novel multivariate model to infer TF regulation of activation-dependent chromatin accessibility (**Fig. 4A, Fig. S5A**). TF motif enrichment analyses nominate potential TF regulators of accessibility patterns, but do not directly leverage important relationships between TF activities and chromatin accessibility signals. Post-hoc assessment of this relationship is non-trivial, when considering the multivariate, context-specific interactions of TFs across the 122 pseudobulk ATAC profiles (as shown in **Fig. 4A**). Thus, we explicitly modeled chromatin accessibility as a function of GRN-derived protein TF activities, using regularized regression to incorporate the TFBS predictions.

Similar to GRN inference, we modeled chromatin accessibility as a multivariate linear function of TF activities, constructing a model for each dynamic accessibility pattern in our dataset:

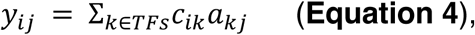

where *y_ij_* ∈ ℝ^|peak^ ^set|×|sample|^ corresponds to pattern i in pseudobulk sample biological replicate j (e.g., “resting Th1, Donor 2”), (*a_kj_* ∈ *R*^|TF|×|sample|^ is the activity of TF k in condition j, as estimated from the final GRN (described above), and *c_ik_* ∈ ℝ^|TF|×|peak^ ^set|^ describes the regulatory effect of TF k on accessibility pattern i. In total, we constructed models for 12 dynamic accessibility patterns, pertaining to the median peak accessibility per cluster (7 RR clusters from **Fig. 4A**, and 5 naïve and memory-shared activation-dependent clusters from **Fig. S5A**).

The prior information was derived from motif-based TFBS prediction in each peak cluster (parameters and motif database as described above). We first counted the fraction of peaks per cluster with at least one binding site and calculated enrichment relative to the background of all reference peaks (one-sided Fisher exact test). For each cluster independently, each TF is ranked according to p-value, yielding adaptive LASSO penalty weighting factors γ*_ik_* ∈ [0,1] in which the most enriched TF k receives the smallest weight and therefore smallest penalty:

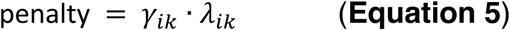

The resulting Λ*_ik_* ∈ ℝ^|peak^ ^set|×|TF|^ penalty matrix is used to guide network inference using an adaptive elastic net objective function:

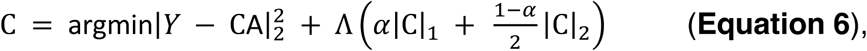

where *Y* ∈ ℝ^|peak^ ^set|×|samples|^ is the accessibility matrix, *A* ∈ ℝ^|TF^*^s^*^|×|samples|^ is the activity matrix, Λ ∈ ℝ^|genes|×|*TFs*|^ is the matrix of penalties associated with each interaction and α ∈ [0,1] is parameter that controls the relative importance of the ridge (L2) versus LASSO (L1) penalty. A small L2 penalty stabilizes the optimization, and we set L = 0.99 to prioritize the L1 penalty and model sparsity(Friedman et al., 2010). Regularized regression was performed with GLMNet v0.7.0 in julia(*JuliaStats*, 2021). *λ_ik_* selection and network interaction confidences were estimated using StARS, as described for GRN inference.

#### Visualization of Accessibility Networks. (e.g., Fig. 4D)

We visualized any regulatory interaction between TF and peak cluster observed in 75 out of 100 subsamples.

### GWAS trait genetic risk variants and chromatin accessibility co-localization analysis

Mirroring the approach in(Wayman et al., 2024), we utilized a curated set of 538 human diseases and phenotypes that had ≥20 independent risk loci from GWAS in European ancestry(Cerezo et al., 2025). We removed the 136 phenotypes associated with lab-collected values, leaving 387 for testing (**Table S8**). We applied the Regulatory Element Locus Intersection (**RELI**) algorithm(Harley et al., 2018) to identify CD4+ T-cell context-specific chromatin accessibility regions that significantly co-localized with genetic risk variant sets (Wayman et al., 2024). Bonferroni correction was applied to account for multiple hypothesis testing. Disease phenotype variant and chromatin accessibility sets were reported as significantly co-localized, if the enrichment achieved P_adj_ < 0.1 and was supported by peak-variant overlaps in at least three independent risk loci. In **Fig. 7A**, phenotypes were clustered using k-means and a distance metric based on Pearson correlation.

### KLF6 ChIP-seq

Whole blood was obtained under IRB2008-0090 at Cincinnati Children’s Hospital Medical Center. PBMCs were isolated using a Ficoll density gradient. Human CD4+ T cells were enriched by negative magnetic selection (Invitrogen; 11346D) and plated at 10 million in 1mL per 6-well flat bottom plate and stimulated with plate bound anti-CD3 (2 µg/mL BE0001-2; Bio X Cell) and soluble anti-CD28 antibodies (2 µg/mL; BE0248; Bio X Cell) in RPMI containing 10% FBS for 5h, then crosslinked and subsequently processed through ChIPmentation, library creation and DNA sequencing as previously described (Yukawa et al., 2020), except that sonication was conducted without SDS in a 1X Tris-EDTA (TE) buffer containing 0.1% NP40 and 0.1% Triton X100.

The ChIP procedure was performed on the Diagenode IP-Star Compact automation system. Antibodies against KLF6 (2µg; 39-6900; Invitrogen or AF3499; R&D systems) were bound to 4.5µL of Protein G Dynabeads and incubated with 2µg of activated CD4^+^ chromatin. Sequential washes with RIPA-150 (buffer 1), RIPA-300 (buffer 2), LiCl (buffer 3) and TE + 0.2% Triton X-100 (buffer 4) ended in the elution in 10mM Tris-HCl containing 0.25M NaCl. Next the complexes were washed with cold 10mM Tris-HCl and incubated with loaded Tn5 transposase (C01070012; Hologic Diagenode) per an established ChIPmentation protocol with an incubation for 20 minutes to accomplish tagmentation of ChIPed DNA (Schmidl et al., 2015). The tagmented ChIPed DNA were washed with RIPA-150, decrosslinked (with proteinase K in TE with 150mM NaCl and 0.3% SDS; incubated at 42°C for 30 minutes, 65°C for 4h, then 15 °C for 10 minutes) and purified from beads using the Qiagen MinElute DNA kit per the manufacturer’s instructions. The library was amplified using a cycle number based on side reaction to stop amplification prior to saturation. The product was cleaned with AMPure beads, and the size and distribution were checked on TapeStation (Agilent). Sequencing was performed using the Illumina NextSeq 2000 at the Genomics, Epigenomics, and Sequencing Core, University of Cincinnati.

Reads were trimmed from the raw fastq files using TrimGalore (v0.6.6)(Felix Kreuger, 2020). Trimmed fastqs were then aligned using Bowtie2 (v2.4.2, additional params: --very-sensitive --maxins = 2000 --B) (Benjamin Langmead, 2020). Aligned bam files were deduplicated using samtools (v1.9) markdup (additional params: -r) and then indexed with samtools. Finally, peak calling was performed with macs2 (v2.1.4, addional params --nomodel --extsize 147 –q 0.01). (Gaspar, 2018). The data passed our quality-control criteria(Cazares et al., 2023b): average read length > 20bp, FASTQC (Andrews S., 2010) report passing for per-base N content, > 500 peak regions (54506 total), and motif enrichment analysis (Heinz et al., 2010)identified the KLF motif family as the second most enriched transcription factor binding motif within the called peaks (-log_10_(P_adj_) = 340).

### Precision-recall evaluation of GRN target genes for KLF6, MAF and PRDM1

#### Gold standard TF-gene interactions

PRDM1 TF ChIP-seq data from CD4 T cells was processed as described above for KLF6 ChIP-seq (Guo et al. 2022). For the human interactions, genes were linked to ChIP-seq peaks by taking all TSSs within 5kb of a peak. Murine “gold-standard” TF-gene interactions for *Prdm1* and *Maf* were derived from (Guo et al., 2022; Wayman et al., 2023) and mapped to human orthologs using biomaRt (v2.54.1)(Durinck et al., 2009). For mouse genes mapping to multiple human genes, all mappings were retained. Following the standard in the field (Marbach et al., 2012), we evaluated precision (fraction of GRN TF-gene interactions supported by KO or ChIP-seq data) and recall (fraction of KO or ChIP-supported interactions recovered by the GRN) as a function of GRN interaction confidence scores. As a point of comparison, we generated GRNs from our multiome data using two other popular methods. In brief, for *CellOracle* (Kamimoto et al., 2023), per their protocol: We generated a base GRN representing potential TF–gene interactions, derived from our snATAC-seq reference peaks. Distal regulatory interactions were inferred based on co-accessibility using *Cicero* (default parameters)(Pliner et al., 2018), while proximal regulatory regions were identified by locating peaks near transcription start sites (TSSs) using *CellOracle*’s HOMER-based motif analysis module (Heinz et al., 2010; Kamimoto et al., 2023). We then integrated the distal (*Cicero*) and proximal (TSS) peaks and retained only those with a co-accessibility score ≥ 0.8. Using the filtered peak set, we performed motif scanning to identify TF-binding motifs using the CIS-BP motif database (v2.0.0)(Weirauch et al., 2014). The resulting TF–gene interaction network served as the base GRN for building cell type-specific GRNS. Following the construction of cell type-specific GRNs, the confidence ranking of regulatory interactions were derived as described in *Kamimoto et al*.: Across subnetworks, we retained regulatory interactions P ≤ 0.001 and interactions were ranked based on the absolute value of the regression coefficients, with p-values used to break ties. Finally, we selected high-confidence TF–gene interactions such that, on average, each gene was regulated by 20 TFs, consistent with *Inferelator* model complexity. For *<u>SCENIC+</u>*: Enhancer gene regulatory networks were inferred using the *SCENIC+* workflow(Bravo González-Blas et al., 2023) with default parameters unless otherwise specified. Chromatin accessibility topic modeling was conducted with pycisTopic using Mallet (150 iterations), testing 11 topic models (2 topics and 5 to 50 in increments of 5). The optimal model, which included 45 topics, was chosen based on *SCENIC+* model selection criteria. Differentially accessible regions between cell types were defined as those with a log_2_(fold-change) > 0.58.

### Analysis of *Soskic et al*. scRNA-seq data

Processed scRNA-seq data (Soskic et al., 2022) were downloaded as a Seurat object from https://trynkalab.sanger.ac.uk. Gene feature Ensembl IDs were converted to human gene symbols using biomaRt (v2.54.1)(Durinck et al., 2009). Ensemble IDs mapping to the same gene were summed. The single-cell expression profiles were aggregated to generate pseudobulk count matrices per donor for each of eight cell type-timepoint combinations (naïve, TCM, TEM or Treg at 0 or 16h), which were filtered for a minimum library size of 30,000. Differential gene expression analysis was performed using the pseudobulk analysis pipeline above (“Pseudobulk batch correction, data normalization and differential analyses”). TFA was estimated from the pseudobulked expression profiles using the GRN constructed from our multiome-seq data (see “*Final transcription factor activity*” in the “GRN analyses” section). For each of the eight cell type-timepoint combinations, we used a two-tailed t-test to assess TFA sex-dependence and Pearson correlation between TFA and donor age to assess age-dependence. P-values were adjusted for multiple hypothesis testing using the Benjamini–Hochberg procedure.

### Analysis of *Monian et al*. scRAN-seq data

Processed scRNA-seq data (Monian et al., 2022)were downloaded as an h5ad file from GEO (GSE158667). The h5ad object was converted to a Seurat object using the *Convert* function in Seurat (v5.1.0). Only cells annotated as high quality from the original publication (HighQ = TRUE) were retained for downstream analysis. Single-cell expression profiles were aggregated per donor for each time point and CD4^+^ T cell population (CD137^+^, CD154^+^ and DN) to generate pseudobulk expression profiles, which were filtered for a minimum library size of 30,000. TFA was estimated using the GRN constructed from our multiome-seq data (see “*Final transcription factor activity*” in the “GRN analyses” section). For each CD4+ T cell population, we treated each donor-timepoint as a biological replicate, reasoning that each timepoint in the peanut oral tolerance study provided insights into distinct T-cell states, based on timing since last antigen encounter and the frequency and number of those encounters proceeding the peanut stimulation timepoint. For example, baseline and avoidance timepoints could be years to decades since last encounter, while build up and maintenance corresponded to recent encounter but clinically important differences in the frequency and number of encounters proceeding stimulation. For **Fig. 6J**, we used two-sided t-tests with Benjamini–Hochberg correction for multiple testing to assess differences in TFA between the three CD4^+^ T cell population (CD137^+^, CD154^+^ and DN).

We compared TF activities of the three CD4+ T cell populations in the *Monian et al.* dataset to simple simulations from our dataset: Assuming a minimum of 5h for antigen processing and presentation for their 20h timepoint (post peanut protein incubation with PBMCs), their CD137+ cells were compared to our activated Treg (average TFA of 5 and 15h timepoints), their CD154+ cells were compared to our activated TEM (average TFA of 5 and 15h timepoints) and their DN cells, assumed to be majority naïve and TCM, were compared to resting TCM and naïve cells (i.e., TF activity estimates were averaged between TCM and naïve cells). Our activated TEM population was constructed by averaging TFA across our five TEM populations (Th1, Th2, Th17, MHCII, CTL). We used ANOVA with Benjamini-Hochberg correction to identify TFs whose activities were not constant across the three groups from both the *Monian et al.* dataset and our simulated groups (CD137^+^, CD154^+^ and DN, P_adj_ < 0.1). We visualized the top 50 most group-dependent (based on P_adj_) from both datasets, resulting in 90 unique TFs. We defined good agreement between TFAs based on their correlation between the two datasets (ρ > 0.5)

### Interactive GRN and multiome-seq data exploration resources

The resources developed in our study have been made interactively available from https://github.com/MiraldiLab/RapidRecall. These resources include: (1) *GRN Visualization:* All subnetworks described can be interacted with through cytoscape v3.10 by loading cytoscape session files (Shannon et al. 2003). (2) *Data Visualization:* We used RShiny (v1.7.4.1) to enable interactive visualization of gene expression, transcription factor activity and data dimension reduction (Chang et al. 2025). (3) *Genome browser of accessibility chromatin and maxATAC TF binding site predicitons*: A track data hub (Raney et al., 2014) was generated to visualize both chromatin accessibility data and the maxATAC predictions in the UCSC Genome Browser (see the “*maxATAC prediction of TFBS*” section). All maxATAC-predicted TFBS were first aggregated into subpopulation-specific BED files across all TFs, resulting in a total of 16 BED files. These BED files were then converted into bigBED format using the *bedToBigBed* utility from version 466 of UCSCtools(Perez et al., 2025), and the data hub was compiled following the format described in the Basic Track Hub Quick Start Guide: (https://genome.ucsc.edu/goldenPath/help/hubQuickStart.html). (4) *Codebases*: The codebases to generate all figures in the manuscript are freely available from the repo as well.

## Acknowledgements

This project was supported by National Institute of Health (**NIH**) funding: R01AI153442 (AB, ERM), U01AI150748 (ERM, LCK), R01AI173314 (ERM, LCK), P30AR070549 (LCK). We thank David A. Hildeman, Matthew T. Weirauch and Yuriy Baglaenko for helpful discussions and advice on the manuscript and the CCRF Single Cell Genomics Facility (RRID:SCR_022653) for assistance optimizing the multiome-seq assay for this study. Some of our images were created with https://BioRender.com (**Fig. 1A**).

## Author contributions

AB and ERM conceived of the study. AB and ERM designed the experiments, which were executed and optimized by SK and SP. AK and ERM designed the computational analyses, which were executed by AK with support from ATB, SMO, ZFY, ANR, JAW and MK. AK, SK, AB and ERM analyzed the experimental data and modeling results. AK, SK, AB and ERM wrote the manuscript. All authors provided feedback on the manuscript.

## Data availability

Raw and processed multiome-seq data, KLF6 ChIP-seq and WGS have been deposited in GEO under accession number GSE282266 (Multiome and KLF6 ChIP-seq) and GSE287942 (WGS).

## Supplemental Figure Legends

**Fig. S1.**
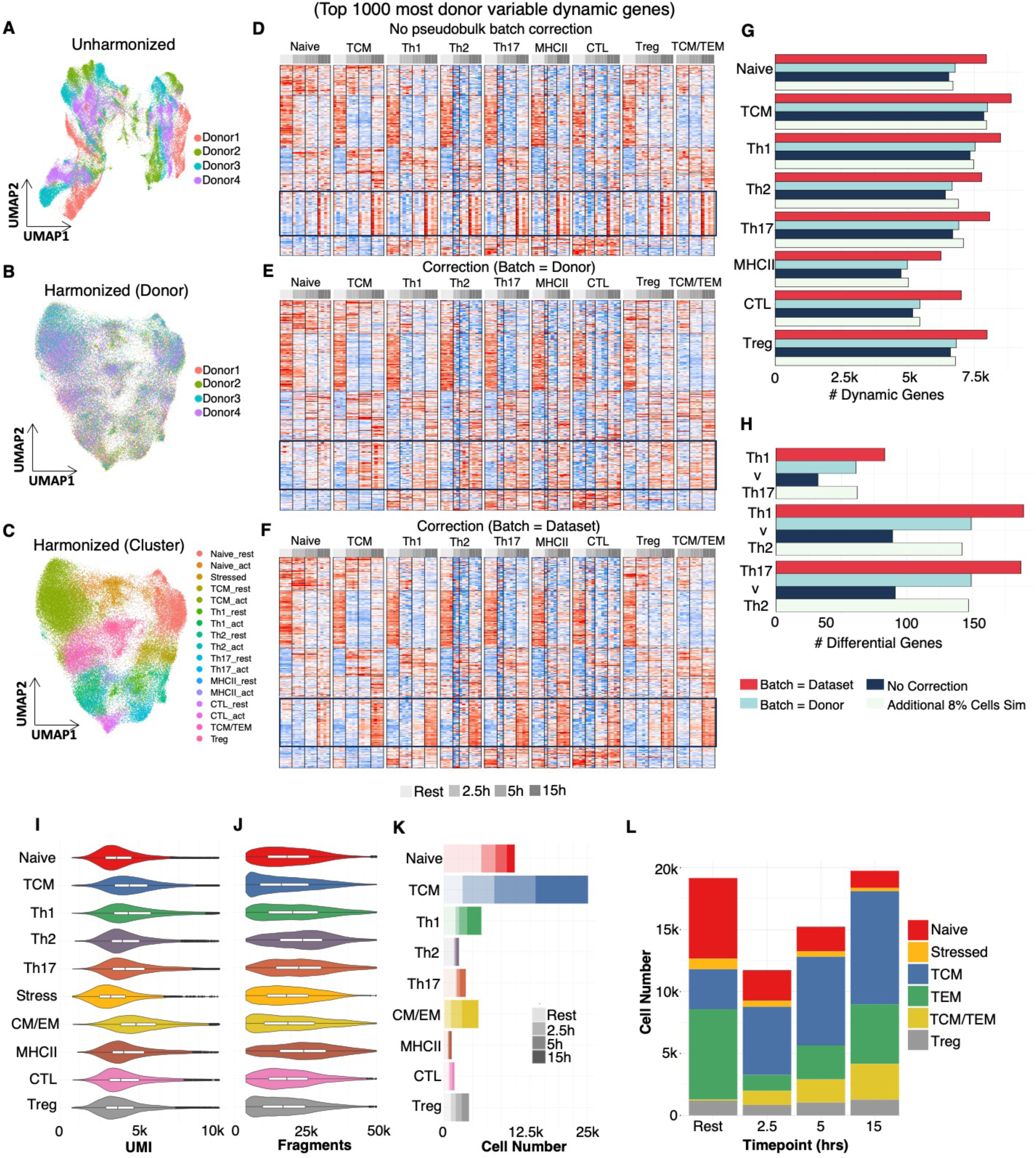
Evaluation of the spike-in donor batch-correction strategy and multiome-seq data characteristics (QC and compositions), related to Fig. 1. UMAP of **(A)** unharmonized and **(B,C)** harmonized data colored by either donor or cell population. Harmony integration reduces the effect of donor on clustering, thereby improving aggregation of nuclei by cell population / timepoint for downstream pseudobulk analyses (see **Methods**). Three batch-correction strategies for pseudobulk expression data were evaluated visually (**D-F**) and quantitatively (**G-H**), see **Methods**. (**D-F**) The 1000 activation-dynamic genes with greatest batch effects are displayed for z-scored VST counts resulting from the three pseudobulk batch-correction strategies: **(D)** no batch correction, **(E)** incorrect “batch = donor” design and **(F)** “batch = dataset” correction enabled by the spike-in strategy. Black box highlights a gene cluster where batch effects are especially apparent in the “no batch” strategy (D) but absent from the other two strategies (E-F). **(G)** Total number of dynamic genes identified by differential DEG analysis for each major T-cell population, using “batch = dataset” (red), batch = donor (light blue) or no batch correction (dark blue). An additional counts matrix was simulated by up-sampling an additional 8% of experimental cells and applying the batch = donor correction (beige) (**Methods**). **(H)** Number of differential genes identified between Th1, Th2 and Th17 cell populations at rest; methods as described for (G). Spike-in correction with batch = dataset resulted in the largest number of differential genes for both comparison types (G,H), providing strong support for using a spike-in control in the experimental design. **(I)** UMI counts and **(J)** fragment counts for each major cell population are relatively consistent. **(K)** Number of cells in each major cell population stacked by timepoint. **(L)** Cell number for each major cell class (Naïve, Stressed, TCM, TEM, TCM/TEM, Treg). TEM is composed of Th1, Th2, Th17, MHCII and CTL.

**Fig. S2.**
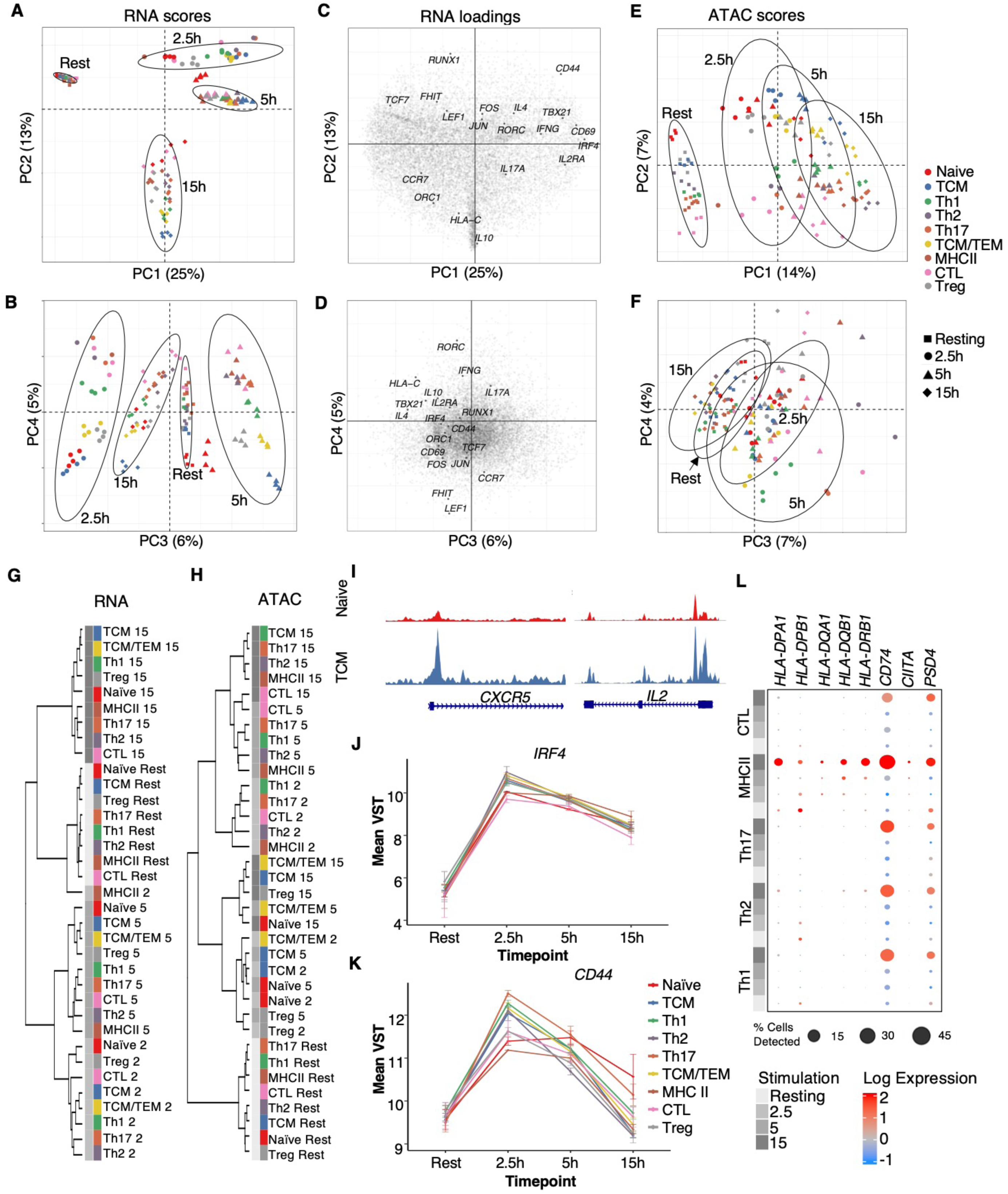
Assessment of variation in the gene expression and chromatin accessibility data, related to Fig. 1. **(A-B)** Principal component analysis (**PCA**) of pseudobulk gene expression profiles. Datapoint color and shape show cell-type and timepoint, respectively. PC1-PC3 correlate with timepoint, while PC4 describes cell-type variation. 95% confidence ellipses are drawn for each of the 4 timepoints. (**C-D**) Loadings plots of selected transcripts contributing to the PCs. **(E-F)** PCA of pseudobulk chromatin accessibility profiles. PC1 describes timepoint variation, while PC2 describes cell-type variation. PC3-4 has a mix of timepoint and cell-type contributions. Hierarchical clustering of the pseudobulk (**G**) expression or (**H**) accessibility profiles. Only features with a coefficient of variation > 0.01 were included. Data were z-scored and clustered using Ward’s method and Euclidean distance. The RNA profiles clustered by timepoint, while clustering of the ATAC profiles reflected a mix of timepoint and cell-type variation. **(I-J)** Normalized expression of activation-inducible *IRF4* and *CD44* in each population. **(K)** Time-dependent expression dotplot of MHCII-associated genes across the TEM populations. Color represents average log expression across the four donors and size represents percent of cells with at least one transcript detected. MHCII cluster has the highest expression of antigen presentation genes among all T cells.

**Fig. S3.**
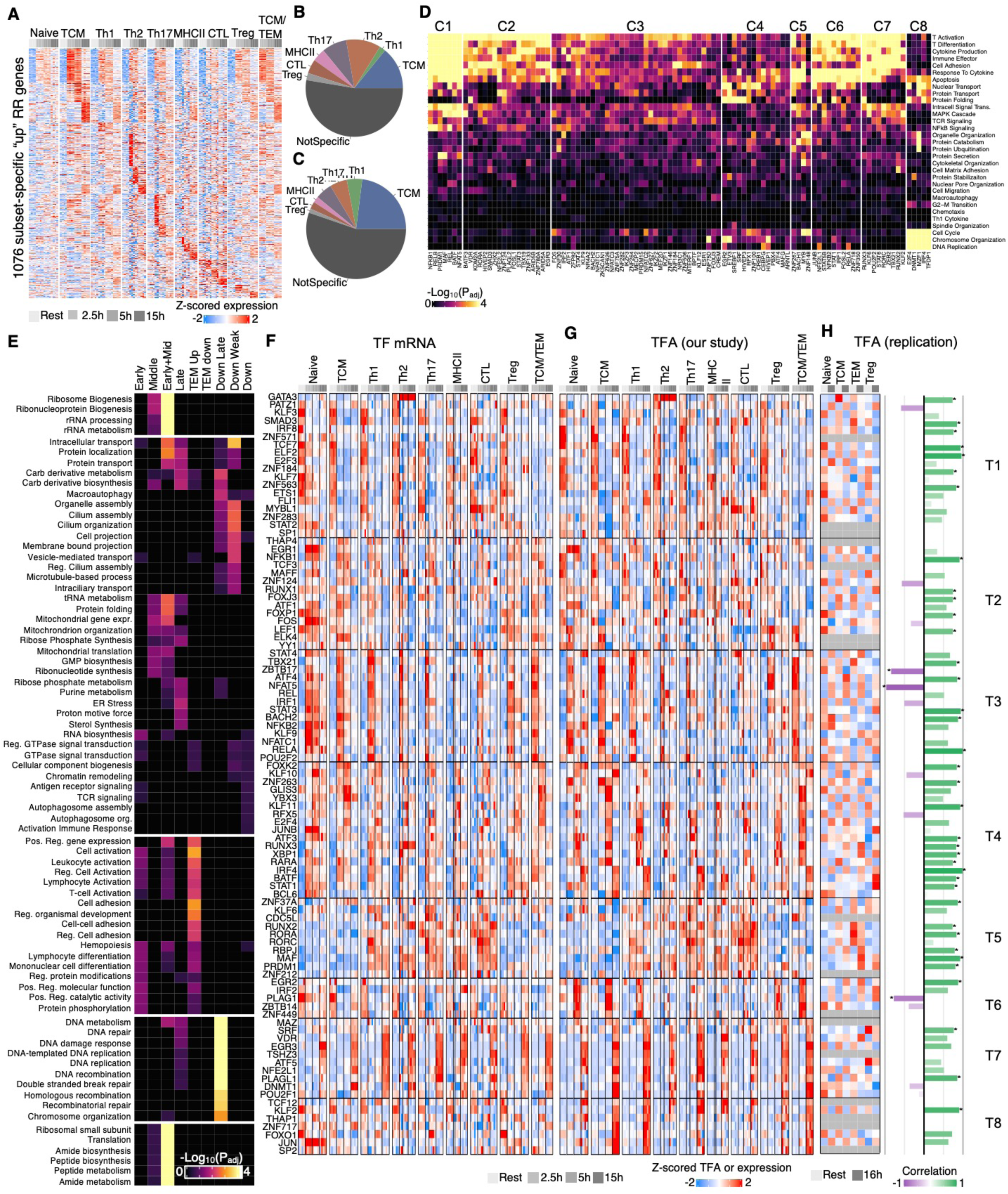
Characterization of T-cell activation transcriptional responses, related to Fig. 2. **(A)** Pseudobulk gene expression heatmap of the 1076 RR genes that are uniquely increased in only one memory population (e.g., “subset-specific”; FDR = 10%, log_2_(FC) > 0.58). The compositions of memory subset-specific-versus non-specific RR genes that are **(B)** up-regulated (1076 out of 2372 genes) or **(C)** down-regulated (1577 out of 3773 genes). Most RR genes (>50%) are defined based on more than one population, and TCM have the greatest fraction of subset-specific RR genes. Compositions are influenced by both biological factors (functional similarity of subsets) and technical, as we had less power to detect differential genes in lower-abundance populations (e.g., CTL and MHCII), **(D)** Pathway enrichments of each TF’s predicted activated target genes, using GO Molecular Pathways. Enriched pathways shown reached significance for at least one TF (Fisher’s exact test, FDR = 10%). Pathway enrichment analysis of each TF’s repressed target genes yielded few significant enrichments and are therefore not shown. **(E)** Pathway enrichment of each RR gene cluster, using GO terms. Any pathway achieving significance in at least one cluster is included (Fisher’s exact test, FDR = 10%). Z-scored **(F)** gene expression and **(G)** TFA resolved by donors, cell types and timepoints, for select TFs (**Methods**). (H) Replication of TFA dynamics using an independent dataset (*Soskic et al*.) for a combination of eight matching cell type-timepoint combinations (naïve, TCM, TEM Treg at 0h or 16h post aCD3/CD28 activation). Grayed boxes denote TFs whose transcripts were not nominally expressed (<5% cells). The adjacent bar plot shows the correlation between TFA profiles across the eight matching cell type conditions between the two datasets (F and G). Asterisk denotes significant correlation (P_adj_ < 0.1).

**Fig. S4.**
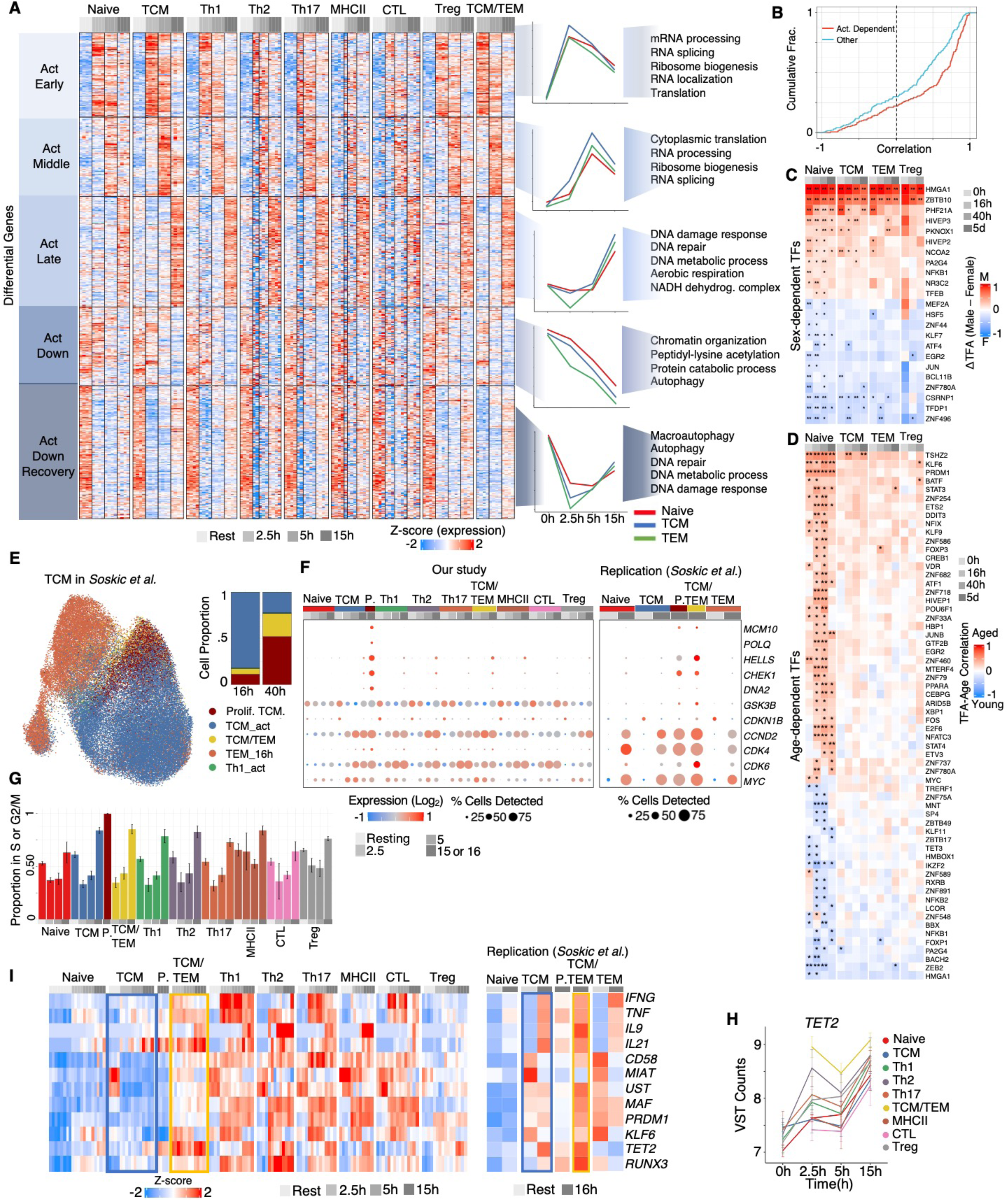
Characterization of shared T-cell activation transcriptional responses, proliferating TCM and TCM/TEM, related to Fig. 2 and Fig. 3. **(A)** K-means clustering of pseudobulk gene expression profiles for all activated, non-RR genes. Gene set enrichment analysis (Fisher’s exact test) with GO terms was performed for each cluster, and the top pathways, achieving FDR = 10%, are listed. Median gene expression for the major T-cell populations is shown to the right of each cluster. TEM is the average of Th1, Th2 and Th17. **(B)** Cumulative distribution function (**CDF**) of Pearson correlation coefficients between TF activity profiles from this study and those from *Soskic et al*. across shared time points (0 h and 15/16 h) and cell types (Naive, TCM, TEM, and Treg) for a total of n = 8 datapoints per TF. For TFs nominally expressed in both datasets, “activation-dependent” TFs (n = 181, red curve) were defined based on differential TFA in our dataset (for at least one cell type (naïve, TCM, TEM, Treg across at least two timepoints) with t-test, P_adj_ < 0.1), while the remainder of nominally expressed TFs compose “other” (n = 197, aqua curve). As activation is the strongest source of transcriptome variation in both datasets, there is greater positive correlation for activation-dependent TFs (K-S test, P < 0.0001). **(C)** The averaged difference in Z-scored TF activities between males and females (Male − Female) per cell type timepoint for TFs exhibiting sex-dependent variation in the *Soskic et al*. dataset (67 males, 52 females, t-test, *P_adj_ < 0.1; ** P_adj_ < 0.01). Positive values indicate higher inferred activity in males (red), and negative values indicate higher activity in females (blue). **(D)** Correlations between TFA and donor age for TFs that are age-dependent in at least one cell type timepoint (e.g., naïve at 0h) in the *Soskic et al*. dataset. Donor ages ranged from 19 to 79 years, with mean age 47, SD 15.6. Asterisks denote significant correlation (*P_adj_ < 0.1; ** P_adj_ < 0.01). **(E)** Label transfer of cell-type annotations from our multiome-seq data to the 16h activated TCM cells in *Soskic et al*. scRNA-seq (left panel). 99% of their 16h TCM cells mapped to our annotations for activated TCM, proliferating TCM or TCM/TEM, and the proportions of proliferating TCM and TCM/TEM were expanded from 16h to 40h TCM timepoints (right panel). **(F)** Select “Proliferating TCM” marker genes, differentially increased in proliferating TCM relative to other T-cell populations at 15h (FDR = 10%, Log_2_(FC) > 0.58), as detected in our dataset (left panel) or as replicated in *Soskic et al.* (right panel). Dot size indicates the percentage of cells with transcript detected, and color represents average expression Log_2_(TPM). **(G)** Percentage of each T-cell population predicted to be proliferating (S, G2/M phase combined). Error bars are standard error (n = 4 donors) per cell-type timepoint. Relative to any other population at any other timepoint, proliferating TCM have a significantly higher proportion of cells in S or G2/M phase (P_adj_ < 0.1). **(H)** Normalized expression of *TET2* across T-cell populations. TCM/TEM has significantly higher expression of *TET2* when compared to all other cell types except Th2 (FDR = 10%, Log_2_FC > 0.58). **(I)** Gene expression of select cytokines and TFs distinguishing TCM/TEM (yellow box) from TCM (blue box), identified in our dataset (left heatmap) and replicated in the *Soskic et al*. dataset (right heatmap).

**Fig. S5.**
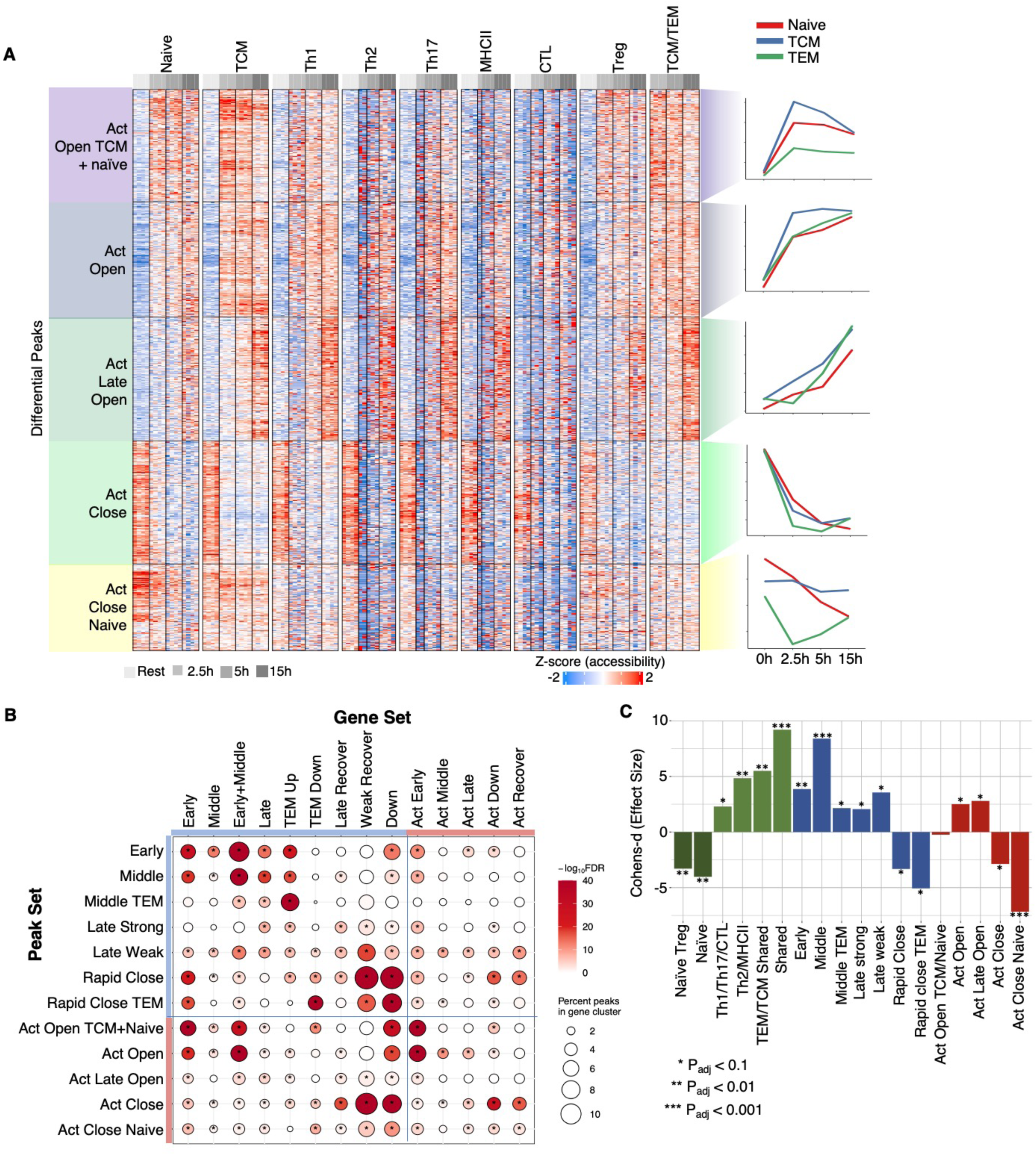
Conserved activation-responsive chromatin accessibility and the enrichment of activation-responsive chromatin in activation-responsive gene sets, related to Fig. 4 and Fig. 6. **(A)** K-means clustering of accessible chromatin regions with shared activation-dynamics across naïve and memory populations. Median peak accessibility for the major T-cell populations are shown to the right of each cluster. TEM is the average of Th1, Th2 and Th17. **(B)** Gene targets for RR (**Fig. 4A)** and memory-naïve shared activation-dependent (A) peak clusters were determined using Trac-looping and Micro-C (when available) or proximity (**Methods**). Fisher exact test was used to identify significant overlap of peak set gene targets and RR (**Fig. 2A**) and naïve-memory shared activation-dependent (**Fig. S4A**) gene clusters. Dot color represents log-transformed adjusted p-values, while the size indicates the percentage of peaks within each peak set linked to genes within each RR gene cluster. **(C)** Cohen’s d effect sizes for module score differences between activated naive at 15h and precocious naive populations for each activation-dependent peak sets (**Figs. 4A, S5A**). Effect sizes were calculated using donor-averaged module scores (n=4 donors per cell type); Cohen’s d is the standardized difference between group means divided by the pooled standard deviation. *P_adj_ < 0.1, **P_adj_ < 0.01, ***P_adj_ < 0.001.

**Fig S6.**
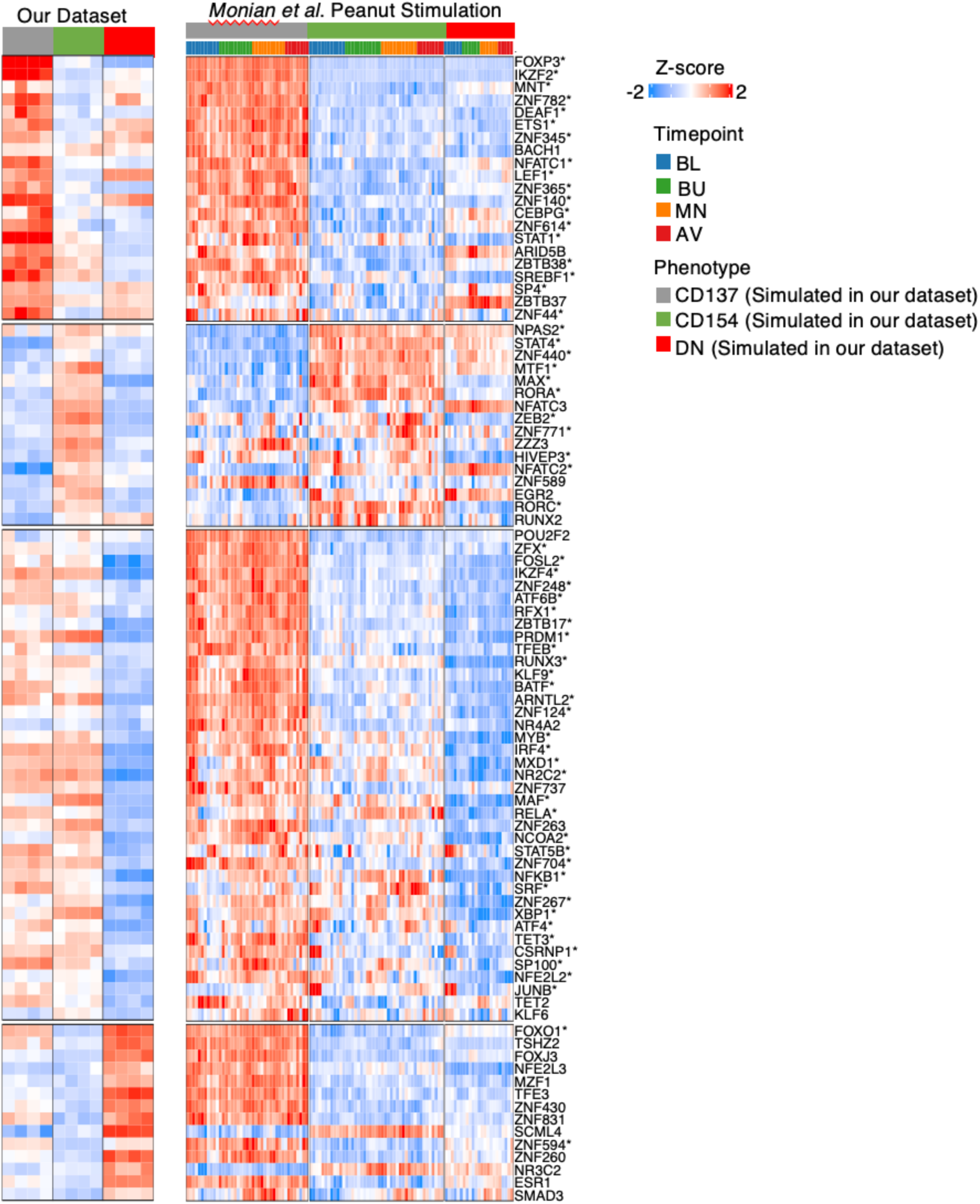
Comparison of TF activities between aCD3/CD28-stimulated and antigen-specific T-cell activation. Comparison of TF activity estimates for populations of CD137^+^, CD154^+^ or double-negative (**DN**) CD4^+^ T cells, as simulated from our aCD3/CD28 T-cell activation data (left panel) or measured from scRNA-seq of sorted peanut-stimulated CD4+ T cells in *Mornan et al*. (right panel), in which cells from peanut-allergic individuals (n = 12) were stimulated *ex vivo* with peanut antigen for 20h, at different timepoints during peanut immunotherapy (BL = baseline, BU = build up, MN = maintenance, AV = avoidance). To approximate the cell populations analyzed by *Monian et al*., we simulated from our dataset: CD137^+^ as the average of Treg cells at 5 and 15 h, and CD154^+^ as the average of TEM subsets (Th1, Th2, Th17, CTL, and MHCII) at 5 and 15 h, DN as the average of naive and TCM cells at 0h. For each dataset, ANOVA was used to identify the top 50 TFs with the most differential activities among the three cell populations, with the union of each set (total of 90 TFs) shown. Asterisks denote “agreement” (ρ_Pearson_ > 0.5) in TFA averages for the three populations (CD137+, CD154+ and DN) between the aCD3/CD28 and antigen-specific T cell activation datasets.

**Fig. S7.**
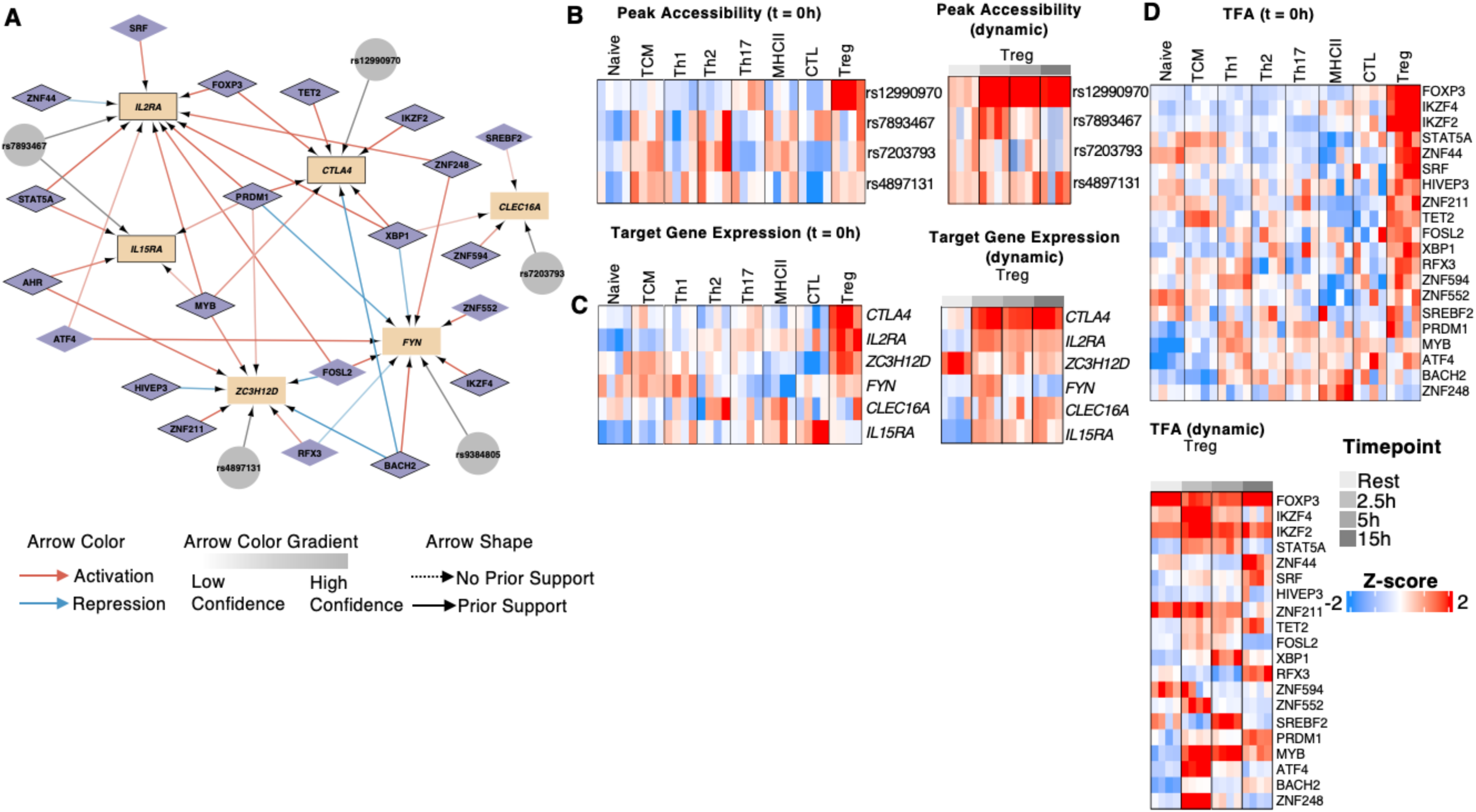
GRN subnetwork for resting Treg chromatin regions co-localized with variants from the “Treg disease” cluster, related to Fig. 7. **(A)** GRNs centered on target genes linked (via trac-loop, micro-C or promoter proximity) to genetic risk variants in the “Treg cluster” (blue box in **Fig. 7A**). Only genes expressed in at least 5% of Treg were included. TF regulators were included if they exhibited Treg-specific activity at rest or post-activation. Outlined nodes represent rapid recall genes for Treg. Red arrows indicate activating interactions, while blue arrows denote inhibitory interactions. Solid arrows represent interactions supported by both gene expression modeling and the ATAC-derived “prior” network, while dashed arrows depict interactions supported by gene expression modeling alone. Arrow color gradient reflects confidence levels, from low to high. (**B**) Peak accessibility (**C**) target gene expression and (**D**) transcription factor activity for TF regulators. For panels B-D, the left heatmap represents the resting state (t=0h), while the right heatmap shows activated dynamics.

## Supplemental Table Legends

**Table S1. Gene regulatory network and functional enrichment of TF target genes.** Predicted TF-gene and TF-peakset interactions from the regulatory network are presented, along with confidence values such as stability scores and the correlation between Transcription Factor Activity (**TFA**) and target gene expression. Additionally, TF functions are annotated through Gene Set Enrichment Analysis (**GSEA**) on each TF’s predicted up-regulated target genes.(**A**) “GRN”, (blue) The list of TF regulator - target genes interactions identified by our GRN ; (**B**) “PeakNetwork”, (yellow) The list of TF regulator - peak set interactions (**C**) “TFFunctionsPositive” (gold) GO Enrichment Analysis Results for targets of each TF regulator with P-adj ≤ 0.1 terms shown (**Fig. S3D**).

**Table S2. Differential gene expression and identification of rapid recall, dynamic, and resting genes.** Differential gene expression results used to identify rapid recall, dynamic, and resting (differential between memory and naïve at baseline) genes. (**A**) “RapidRecallGenes” (blue) Genes categorized as rapid recall (both up-and down-regulated during activation) for any cell type (**Fig. 2A**). (**B**) “ActGenes” (green) genes that are differential between resting and activated (2.5h, 5h, 15h) timepoints (**Fig. S4A**). (**C**) “RestingGenes” (yellow) Genes that are differential between naive and memory populations for the timepoint 0h (**Fig. 5E**).

**Table S3. Rapid recall and dynamic gene cluster enrichments.** We performed GSEA on each rapid call (**Fig. 2A**) and dynamic (**Fig. S4A)** gene cluster to identify functional roles. Similarly, we preformed enrichment of predicted TF up-regulated target genes in each cluster to find likely TF regulators. (**A**) “RR_TFEnrichment” (blue). The list of TFs that are predicted to regulate genes in each RR cluster in **Fig. 2A**. (**B**) “RR_PathwayEnrichment” (green) GO analysis results for each cluster in **Fig. 2A**. (**C**) “Act_TFEnrichment” (yellow) provides the list of TFs that are predicted to regulate genes in each activation dynamic cluster in **Fig. S4A**. (**D**) “Act_PathwatEnrichment” (purple). GO analysis results of each cluster from **Fig. S4A**. P-adj ≤ 0.1 terms are shown.

**Table S4. Enrichment of late (15h) rapid recall genes in cell-cycle related GO pathways.** GSEA results for each memory population’s 15h rapid recall genes for cell-cycle related pathways in the GO database. “TCM/TEM” was not included because this cell population does not have a baseline (t=0) to compare to.

**Table S5. Differential peak accessibility and identification of rapid recall, dynamic, and resting peaks.** Differential peak accessibility results used to identify rapid recall, dynamic, and resting (differential between memory and naïve at baseline) chromatin regions. We also include the cluster identity of each peak from **Fig. 4A**, **Fig. S5A**, and **Fig. 5A**. (A) “RapidRecallPeaks” (blue) Peaks categorized as rapid recall (both up-and down-regulated during activation) for any cell type (**Fig. 4A**). (B) “ActPeaks”(green) Peaks that are differential between Resting and Activated (2.5h, 5h, 15h) timepoints (**Fig. S5A**). (C) “RestingPeaks” (yellow) Peaks that are differential between naive and memory populations for the timepoint 0h (**Fig. 5A**).

**Table S6. TF motif enrichment in peak clusters.** We calculated the enrichment of each TF’s binding motifs from the CIS-BP database in the rapid recall (**Fig. 4A**), dynamic (**Fig. S5A**) and resting (**Fig. 5A**) peak sets.

**Table S7. Differential gene expression and gene accessibility between naïve (t=15h) and naïve precocious.** DESeq2 results for the comparison between activated naïve (t=15h) and the naïve precocious population. This table includes results for gene expression (A, blue) and chromatin accessibility (B, green) (total accessibility across promoter and gene body) (**Fig. 6G**).

**Table S8. Regulatory Element Locus Intersection (RELI) enrichments.** Enrichment of peak-locus intersections for each phenotype across the rapid recall, dynamic, and resting peaksets. The table includes number of overlaps and statistics associated with the enrichment for each condition.

**Table S9. Peak to gene linkages from proximity, Trac, and micro-C.** We linked each peak to a gene based on proximity (+/- 1kb) (**A**), Trac-looping (**B**), and micro-C (**C**). This table includes each peak-gene association separated by linkage method. Trac-loop data are for resting and 16 hrs activated total human CD4 T cells (Lai et al, 2018). Micro-C data are for resting and 24 hrs activated total human CD4 T cells (Oguchi et al., 2024).

